# Beyond the Canonical HRF: Flexible Temporal Modeling Reveals Unconstrained BOLD Profiles During Naturalistic Viewing

**DOI:** 10.1101/2025.11.07.687226

**Authors:** Xin Di, George B. Hanna, Bharat B. Biswal

## Abstract

Naturalistic stimuli, such as movies and narratives, are increasingly used in cognitive neuroscience to map cognitive and affective processes onto brain activity measured with functional MRI (fMRI). Features extracted from movies span multiple levels, from computational visual and auditory inputs to physiological signals and subjective ratings. However, the temporal alignment between these features and the blood-oxygen-level-dependent (BOLD) response varies considerably, and the commonly used canonical hemodynamic response function (HRF) with temporal derivatives may not adequately capture these delays. In this study, we analyzed three movie-watching datasets using cross-correlation and finite impulse response (FIR) deconvolution to map the unconstrained temporal dynamics of visual, auditory, pupillary, and Theory of Mind (ToM) features across the brain. Our results demonstrate that while the canonical HRF effectively captures basic sensory features, it introduces systematic misalignments for inherently delayed signals. Because physiological markers (pupil size) and subject reports (ToM) intrinsically lag the underlying neural events, standard HRF convolution overcompensates for their biological latency, introducing a redundant phase mismatch or "double-delay." Furthermore, our unconstrained FIR models revealed distinct inter-regional temporal hierarchies across the cortex. Recognizing the inherent collinearity of real-world stimuli, these estimated profiles capture the bundled, multi-dimensional dynamics of naturalistic processing rather than perfectly isolated feature effects. Ultimately, these findings highlight the necessity of flexible, reliability-tested temporal modeling to accurately map the complex processing timescales engaged during naturalistic viewing.

## 1. Introduction

Naturalistic stimuli — such as movies or spoken narratives — have become increasingly popular in functional MRI (fMRI) research because they offer tools to investigate brain function under ecologically valid conditions (Di et al., 2026; Hasson et al., 2004; Vanderwal et al., 2019). Unlike traditional, highly controlled experimental paradigms, naturalistic stimuli capture the richness and temporal continuity of real-world experiences, thereby engaging a wide range of perceptual, cognitive, and affective processes simultaneously. This shift has opened new avenues for studying how the brain integrates complex, dynamic information over time.

A seminal line of work demonstrated that when multiple individuals watch the same movie or listen to the same story, their brain activity exhibits significant inter-subject correlations (ISC), revealing shared neural responses to naturalistic events (Hasson et al., 2004). Such data-driven analyses have been instrumental in identifying brain regions involved in high-level perception, attention, and social cognition (Kauppi et al., 2010). However, while these approaches effectively capture synchronized patterns of activity across individuals, they do not directly reveal which specific features of the stimulus drive these neural responses. To address this, researchers have increasingly focused on extracting quantitative features from naturalistic stimuli and mapping their temporal dynamics onto brain activity measured by fMRI (Bartels et al., 2008; Bartels and Zeki, 2004; Huth et al., 2016; Lahnakoski et al., 2012a; Raz et al., 2016).

A wide variety of stimulus-derived features can be used for such mapping. These range from low-level sensory features — such as visual luminance, contrast, and auditory pitch — to higher-level variables derived from physiological signals (e.g., pupil size) or subjective behavioral ratings (e.g., perceived emotional valence or theory of mind, ToM). In recent years, feature extraction has also benefited from advances in deep neural networks, where model-based representations capture complex visual, auditory, or semantic content (McMahon et al., 2023; Samara et al., 2025; Sohn et al., 2024). These features can be analyzed individually using general linear models (GLMs) or jointly in encoding-model frameworks to predict regional brain activity.

Crucially, naturalistic stimuli are not monolithic; they contain deeply nested, hierarchical temporal structures. For example, visual streams progress from rapid, high-frequency shifts in luminance to slower transitions between objects and scenes, while auditory streams build from transient phonemes to extended narratives. A prominent theoretical framework posits that the brain processes these nested features through a topography of hierarchical cortical timescales, or temporal receptive windows (TRWs) (Lerner et al., 2011; Meer et al., 2020). In this framework, primary sensory regions possess short TRWs to process rapid physical properties, whereas higher-order associative networks possess long TRWs to integrate semantic and narrative information over extended durations.

Beyond extrinsic stimulus features, naturalistic viewing also engages internally generated, observer-dependent dynamics. Physiological responses, such as pupil size, act as markers of autonomic arousal and cognitive load, reflecting an integration of sensory input and internal state (Joshi and Gold, 2020). Similarly, subjective behavioral ratings—such as ToM annotations—capture conscious, high-level social-cognitive evaluations of characters’ mental states over time (Saarimäki, 2021). As illustrated in our conceptual framework (Figure 1), appropriately mapping naturalistic viewing to brain activity requires integrating three distinct temporal levels: 1) the nested intrinsic hierarchies of the stimulus features; 2) the biological and cognitive integration delays inherent to the observer’s physiological and behavioral responses; and 3) the region-dependent hemodynamic brain responses.

**Figure 1.**
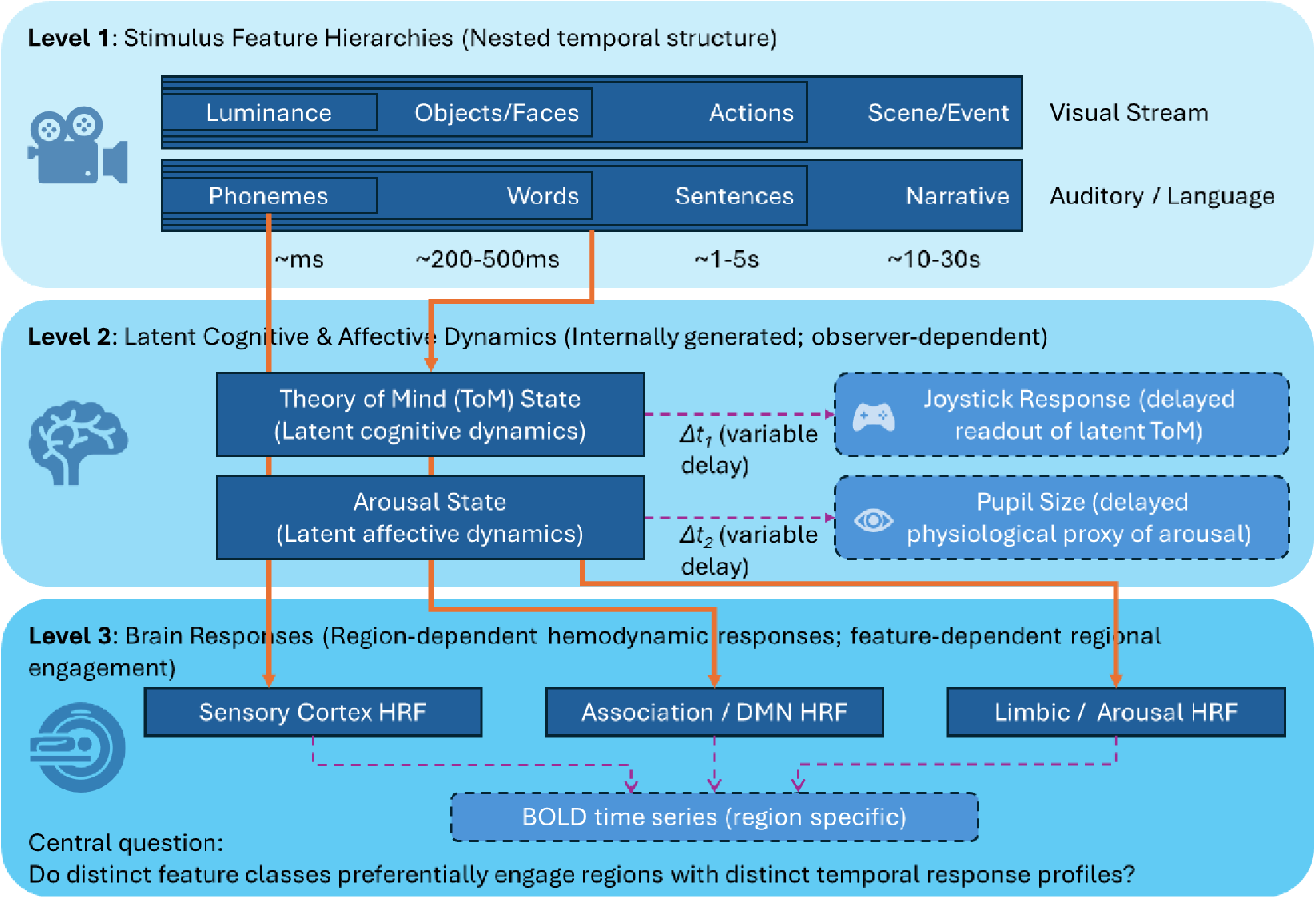
Conceptual framework of nested temporal hierarchies and their mapping to brain responses during naturalistic viewing. **Level 1: Intrinsic stimulus hierarchies.** Naturalistic movies possess a deeply nested temporal structure across modalities. Visual features (e.g., luminance, objects, actions, scenes) and auditory/language features (e.g., pitch, phonemes, words, narratives) unfold over timescales ranging from milliseconds to tens of seconds. The brain processes these continuous streams through a topography of temporal receptive windows (TRWs), ranging from short integration windows in primary sensory areas to extended timescales in higher-order networks. **Level 2: Observer-dependent delays.** Extrinsic stimulus features elicit internally generated latent cognitive and affective dynamics, which are not directly observable. Higher-order states, such as Theory of Mind (ToM) engagement and autonomic arousal, evolve continuously and are captured via delayed behavioral (joystick ratings) and physiological (pupil size) readouts. The integration and motor/autonomic delays between the underlying latent neural states and these observable proxies (Δt₁, Δt₂) introduce critical temporal misalignments. **Level 3: Feature-associated BOLD dynamics.** These multi-level features engage distinct brain regions, each possessing specific neurovascular coupling properties. Because different feature classes preferentially recruit different neural systems (e.g., sensory, associative/Default Mode, and limbic networks) and operate on different intrinsic timescales, the resulting BOLD signals exhibit highly diverse temporal response profiles. Consequently, mapping these features to fMRI requires flexible temporal modeling, as applying a uniform canonical hemodynamic response function (HRF) to the already-delayed readouts from Level 2 risks compounding delays. *Note:* Solid arrows indicate intrinsic neural propagation and hypothesized pathways linking stimulus features to latent cognitive states and regional brain responses. Dashed arrows indicate the delayed measurement of these latent states via observable behavioral or physiological proxies. Δt denotes the variable temporal delays inherent to these readouts.

A critical challenge in linking these features to fMRI data lies in the temporal mismatch between stimulus features and the hemodynamic response. The fMRI signal—via the blood-oxygen-level-dependent (BOLD) effect—typically lags behind underlying neural activity by about 5–6 seconds (Friston et al., 1998; Handwerker et al., 2004). To account for this, most studies either apply a fixed delay (e.g., 4 s) or convolve the feature time-series with a canonical hemodynamic response function (HRF). Some approaches further include temporal derivatives of the HRF to allow limited flexibility in response timing (i.e., ≈1 s adjustment). However, the shape and latency of the HRF vary considerably across brain regions, across individuals, and even within the same region under different physiological conditions (Aguirre et al., 1998; Handwerker et al., 2004). Such intrinsic biological variability complicates efforts to accurately align continuous stimulus features with corresponding neural activity.

This temporal uncertainty becomes especially pronounced when analyzing multiple types of features that occupy different levels of the naturalistic processing hierarchy. Applying a uniform canonical HRF across all feature classes fundamentally ignores the diverse intrinsic frequencies and delays inherent to these distinct signals. Fast sensory inputs, delayed physiological signals, and slow cognitive annotations possess vastly different temporal relationships to neural and vascular processes. Because pupil size and subjective behavioral ratings are themselves delayed responses relative to the eliciting neural activity, convolving them with a standard HRF risks compounding delays. This conventional approach introduces redundant temporal shifts, or "double-delay," that slightly misaligns with the absolute peak of neurovascular coupling (Lloyd et al., 2023).

To overcome the limitations of the canonical HRF and accurately map hierarchical processing timescales, more flexible temporal modeling is required. Finite Impulse Response (FIR) deconvolution provides a robust, non-parametric alternative (Glover, 1999). Rather than imposing a predetermined temporal shape and fixed latency, FIR models utilize a set of time-lagged predictors to directly estimate the unconstrained, feature-associated system response function from the continuous fMRI data. By allowing the response shape and peak latency to vary freely, FIR deconvolution can capture the diverse temporal profiles of naturalistic processing—ranging from rapid sensory responses to the complex, multiphasic interactions of peripheral physiology, and the extended, diffuse hemodynamic delays that characterize slower, higher-order cognitive features.

In the present study, we analyzed three movie-watching fMRI datasets to systematically characterize how different classes of stimulus, physiological, and behavioral features are expressed in their unconstrained BOLD response shapes. We extracted continuous features spanning the hierarchical levels of naturalistic processing: low-level sensory features (frame luminance, frame contrast, and auditory pitch), a physiological measure (pupil size), and a subjective behavioral measure (ToM ratings). Moving beyond conventional cross-correlation analyses, we applied formal FIR deconvolution to estimate the BOLD system response functions across the whole brain. We hypothesize that these diverse features will exhibit distinct response profiles that systematically deviate from the canonical HRF. By mapping these unconstrained temporal trajectories and peak latencies and evaluating their cross-cohort reliability, we aim to empirically reveal the underlying inter-regional temporal hierarchies involved in naturalistic viewing.

## 2. Materials and Methods

### 2.1. Movie-watching fMRI Data

We analyzed three movie-watching fMRI datasets previously used in our work (Di et al., 2023; Di and Biswal, 2022): two from the Healthy Brain Network (HBN) project (Alexander et al., 2017), and one from the Partly Cloudy dataset (Richardson et al., 2018).

#### 2.1.1. Healthy Brain Network (HBN) Dataset

The HBN dataset was obtained from the project repository (http://fcon_1000.projects.nitrc.org/indi/cmi_healthy_brain_network/). For the current study, we included a subset of participants without diagnosed psychiatric disorders who underwent fMRI scanning while watching two video clips: *The Present* (3 min 21 s; Filmakademie Baden-Wuerttemberg, 2014) and a 10-minute segment from *Despicable Me* (Illumination, 2010).

In total, 87 participants watched *The Present* and 83 watched *Despicable Me*, with 66 participants overlapping, yielding 104 unique individuals (61 males, 43 females; age range = 5.0–21.9 years, mean = 12.0 ± 4.1). Additional inclusion and exclusion details are provided in Di et al. (2023).

MRI data were collected across two imaging centers—Rutgers University Brain Imaging Center (RUBIC) using a 3T Siemens Trio scanner, and the Citigroup Biomedical Imaging Center (CBIC) using 3T Siemens Prisma scanners. Scanning protocols were harmonized across sites. Functional MRI parameters were: TR = 800 ms, TE = 30 ms, flip angle = 31°, voxel size = 2.4 × 2.4 × 2.4 mm³, and multiband acceleration factor = 6. Each run contained 250 volumes for *The Present* and 750 volumes for *Despicable Me*. High-resolution T1-weighted anatomical scans were acquired using either Human Connectome Project (HCP) or Adolescent Brain Cognitive Development (ABCD) protocols and were used solely for preprocessing. Additional acquisition details can be found on the HBN website and in Alexander et al. (2017).

#### 2.1.2. Partly Cloudy Dataset

The *Partly Cloudy* dataset was accessed from OpenNeuro (https://openneuro.org/datasets/ds000228; Richardson et al., 2018). It included two groups: adults (n = 29; 17 females, 12 males; age range = 18–39 years, mean = 24.6 ± 5.3) and children (n = 53; 28 females, 25 males; age range = 3.5–12.3 years, mean = 7.0 ± 2.5).

Participants watched a silent version of the Pixar short film *Partly Cloudy* (5.6 minutes; https://www.pixar.com/partly-cloudy) during scanning on a 3T Siemens Tim Trio system.

Younger children were scanned using one of two 32-channel custom head coils, while older children and adults used the standard Siemens 32-channel coil. Functional images were acquired using a gradient-echo EPI sequence (TR = 2000 ms, TE = 30 ms, flip angle = 90°) with 32 interleaved near-axial slices (EPI factor = 64).

Voxel sizes and slice gaps varied slightly across subgroups: (a) 3.13 mm isotropic, no gap; (b) 3.13 mm isotropic, 10% gap; (c) 3 mm isotropic, 20% gap; and (d) 3 mm isotropic, 10% gap. All functional images were resampled to 3 mm isotropic voxels during preprocessing. Each run contained 168 volumes, with the first four discarded to allow for steady-state magnetization. T1-weighted anatomical images were collected in 176 interleaved sagittal slices (1 mm isotropic; GRAPPA = 3; FOV = 256 mm). Further methodological details are provided in Richardson et al. (2018).

### 2.2. Eye tracking data

Although eye-tracking data were not collected during the MRI sessions, a subset of participants from the HBN project participated in a separate EEG session conducted on a different day outside the scanner (Langer et al., 2017). During this session, participants watched several videos, including *The Present*, a shorter segment of *Despicable Me*, *Fun with Fractals* (an educational video), and a trailer for *Diary of a Wimpy Kid*, while simultaneous EEG and eye-tracking data were recorded. Eye position and pupil size were measured using an infrared video-based eye tracker (iView-X Red-m, SensoMotoric Instruments GmbH) operating at sampling rates of 30, 60, or 120 Hz. In the current study, only pupil size data were analyzed. All available recordings were included, but participants were excluded if more than 10% of time points were missing, typically due to blinks or signal loss. The final sample consisted of 1,884 participants (1,032 males, 582 females, 270 not reported), ranging in age from 5 to 21 years (mean = 10.26, SD = 3.38, 270 not reported).

### 2.3. Subjective Rating Data Collection

Subjective rating data were collected from a separate group of participants at the New Jersey Institute of Technology (NJIT) to capture continuous, moment-to-moment fluctuations in perceived mental states of movie characters. A total of 34 participants (age range: 18–27 years, mean ± SD = 20.9 ± 2.3 years; 17 female) took part in the study. All participants reported normal or corrected-to-normal vision and hearing, and none had a history of neurological or psychiatric disorders. Informed consent was obtained from all participants in accordance with the Institutional Review Board (IRB) of NJIT.

Participants viewed the same three video clips used in the fMRI experiment. Each session took place in a quiet, darkened room to minimize visual and auditory distractions. Stimuli were presented on a computer monitor with synchronized sound through a speaker.

Continuous subjective ratings were acquired using CARMA software (Continuous Affect Rating and Media Annotation; (Girard, 2014) with a gaming joystick as the input device.

Participants were instructed to continuously indicate the extent to which they could perceive the characters’ emotions, thoughts, or intentions while watching each clip. This measure was designed to capture participants’ theory-of-mind engagement—that is, the degree to which they inferred or empathically understood the internal mental states of the characters throughout the narrative. Before the experiment, participants completed a short practice session to familiarize themselves with the rating procedure.

The joystick’s vertical position corresponded to a continuous numerical scale ranging from -10 (not at all) to 10 (very strongly), representing the perceived strength of mental-state understanding at each time point. Rating data were sampled at 4 Hz (every 250 ms) to provide a high-resolution temporal record of evolving subjective experience. Between clips, participants were given brief rest periods to minimize fatigue.

The finalized rating data have been uploaded to the Open Science Framework (OSF) repository and will be made publicly available upon acceptance of the manuscript: https://osf.io/qky8p/overview?view_only=f31e53525b514ecaa91a7a8e6e113cb7.

### 2.4. Video feature extraction

We extracted three sensory features from the videos: mean pixel intensity, standard deviation of pixel intensity, and pitch. For the two visual features, each video frame was first converted to grayscale, and the mean and standard deviation of pixel intensity across the two-dimensional matrix were computed. These yielded two time series corresponding to frame luminance and frame contrast, respectively. For the auditory feature, audio signals from all channels were averaged, and pitch was computed using a 100-ms window with no overlap, resulting in a 10 Hz pitch time series. Pitch was extracted only for *The Present* and *Despicable Me*, as the *Partly Cloudy* video contains no sound. Figure 2 shows the time series of mean and standard deviations of pixel intensity and pitch for the video *The Present*.

**Figure 2.**
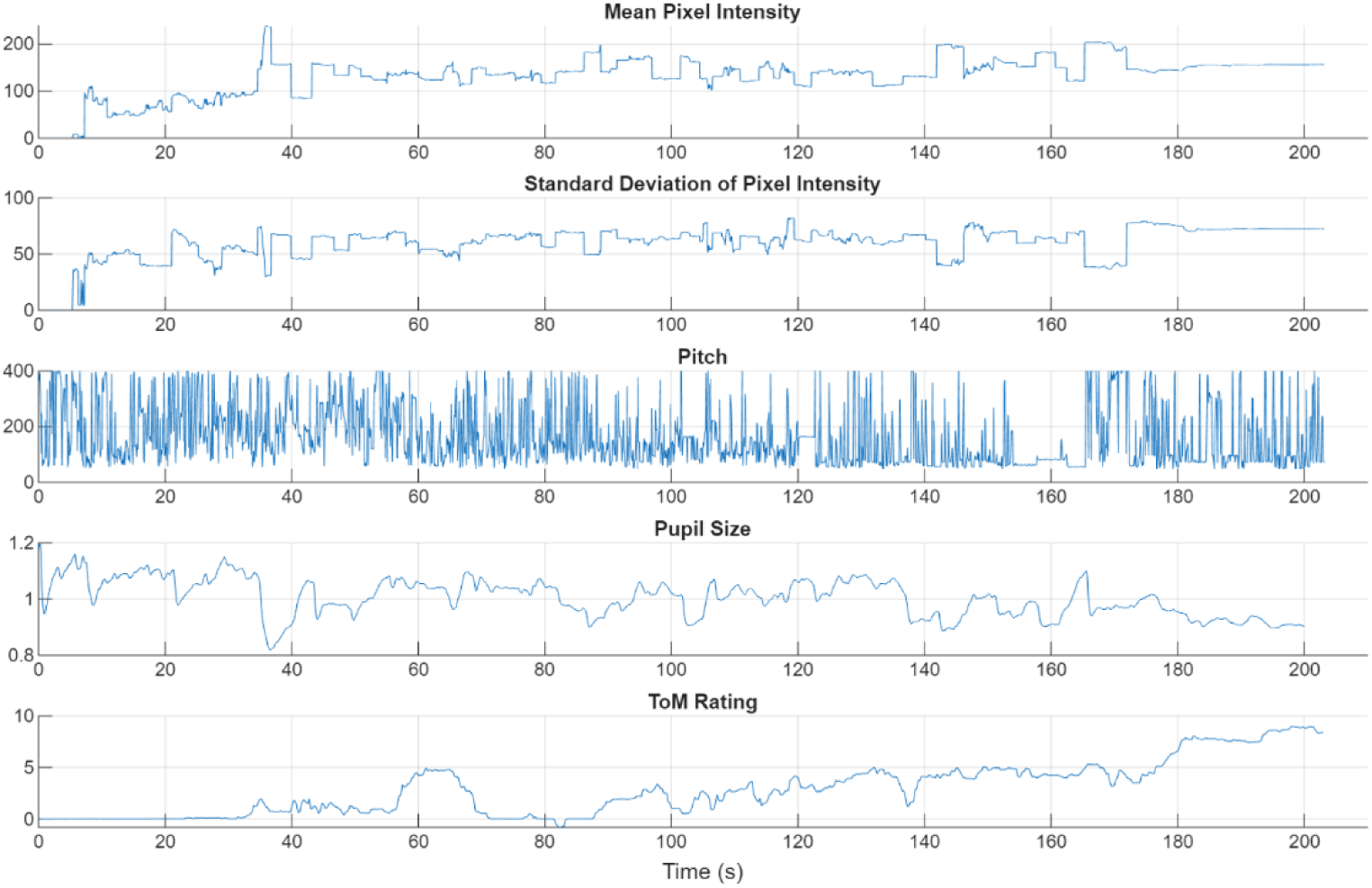
Continuous time series of multimodal features associated with the movie *The Present*. The waveforms illustrate the diverse macroscopic temporal properties across the five feature classes. Low-level sensory features (frame luminance, frame contrast, and auditory pitch) exhibit rapid, high-frequency fluctuations directly tied to instantaneous stimulus changes. In contrast, the physiological (pupil size) and subjective cognitive (Theory of Mind [ToM] rating) measures operate on noticeably slower, lower-frequency timescales, reflecting underlying biological and cognitive integration delays.

### 2.5. Pupil Size Data Processing

Pupil size data for each participant were first resampled to a uniform sampling rate of 30 Hz. Missing values (NaNs) were interpolated linearly, and blink periods—identified as time points with zero values—were also replaced using linear interpolation. Participants with more than 10% of time points classified as blinks were excluded from further analyses. To reduce inter-individual variability in absolute pupil size, each participant’s pupil time series was normalized by dividing by its mean.

To evaluate the consistency of pupil size fluctuations across participants, we computed leave-one-out inter-individual correlations. For each participant, their pupil size time series was correlated with the mean time series of all other participants, resulting in one correlation coefficient per individual. The statistical significance of the group-level mean correlation was assessed using a circular time-shift randomization procedure with 1,000 iterations to generate a null distribution (Kauppi et al., 2010). Finally, the median pupil size time series across participants was extracted and used as the representative group-level signal for subsequent analyses (Figure 2).

### 2.6. Rating Data Processing

The subjective rating data were analyzed using the same leave-one-out inter-individual correlation approach described above. For each video clip, the median rating time series across participants was computed and used as the group-level representative signal for further analyses (Figure 2 shows an example).

### 2.7. Temporal characteristics of multimodal features

Across the five selected features, we observed distinct temporal and frequency characteristics that correspond to their respective levels in the naturalistic processing hierarchy. Low-level sensory features, such as frame luminance and auditory pitch, exhibited highly transient, high-frequency fluctuations (sampled at video frame rates or 10 Hz, respectively) reflecting rapid, instantaneous changes in the external physical stimulus. In contrast, frame contrast exhibited slightly slower dynamic shifts, corresponding to sustained visual scene characteristics. Finally, the physiological (median pupil size) and behavioral (median ToM rating) features operated on markedly slower, lower-frequency timescales. These differences highlight that higher-level features inherently incorporate sluggish biological constraints (e.g., autonomic nervous system delays in pupillary constriction) and higher-order cognitive integration over extended narrative windows. Recognizing these diverse intrinsic frequency profiles is critical, as applying a uniform, fixed-delay HRF to all features risks severe temporal misalignment, directly motivating our subsequent use of flexible FIR deconvolution.

### 2.8. MRI Data Processing

Preprocessing of the fMRI data was performed separately for the HBN and *Partly Cloudy* datasets, following procedures described in previous studies. Here, we provide a brief overview. Structural MRI images for each participant were first segmented and normalized to the standard Montreal Neurological Institute (MNI) space. For *The Present* and *Despicable Me* datasets, the first 10 functional volumes were discarded, resulting in 240 and 740 volumes, respectively. In contrast, all functional volumes were retained for the Partly Cloudy dataset. The remaining functional images were then realigned to the first retained image, coregistered to the participant’s structural image, normalized to MNI space using the deformation fields obtained during segmentation, and spatially smoothed with an 8 mm full-width-at-half-maximum (FWHM) Gaussian kernel.

Finally, a general linear model (GLM) was constructed to regress out head motion effects using Friston’s 24-parameter motion model, which also incorporated an implicit high-pass filter with a cutoff frequency of 1/128 Hz. The resulting residual time series were used in all subsequent analyses.

We adopted a region-of-interest (ROI)–based approach to achieve a balance between spatial specificity and temporal flexibility. Cortical regions were defined using the Schaefer 100-region parcellation (Schaefer et al., 2018), and subcortical regions were defined using the Automated Anatomical Labeling (AAL) atlas (Tzourio-Mazoyer et al., 2002). Fourteen bilateral subcortical nuclei were included: hippocampus, parahippocampus, amygdala, caudate, putamen, pallidum, and thalamus. The Schaefer parcellation was divided into left (#1–50) and right (#51–100) hemispheres and further organized into seven large-scale functional networks: visual, somatomotor, dorsal attention, salience/ventral attention, limbic, control, and default mode.

In total, 114 ROIs were analyzed. For each preprocessed fMRI run, the mean time series was extracted from each ROI, resulting in data matrices of size 240 × 114 for *The Present*, 740 × 114 for *Despicable Me*, and 168 × 114 for *Partly Cloudy* for each participant.

### 2.9. Cross-correlation Analysis

Each feature time series from the three movie clips was first convolved with the canonical HRF implemented in SPM12 (using the default parameters: peak delay = 6 s, undershoot delay = 16 s, and ratio = 6). The convolved feature time series were then resampled to match the temporal resolution of the corresponding fMRI data (TR = 0.8 s for *The Present* and *Despicable Me*; TR = 2.0 s for *Partly Cloudy*).

Subsequently, cross-correlation analyses were performed between the HRF-convolved feature time series and the mean fMRI time series from each ROI for each participant.

Correlations were computed across a lag range of ±16 seconds (i.e., ±20 TRs for HBN data and ±8 TRs for *Partly Cloudy*), resulting in a lag-by-ROI correlation matrix for each participant. For visualization and group-level interpretation, these matrices were averaged across participants after Fisher’s *z* transformation of the correlation coefficients.

In addition, a regression analysis was conducted using the HRF-convolved feature time series as predictors in a general linear model (GLM) framework for each participant. Group-level statistical significance of the resulting parameter estimates was assessed using one-sample *t*-tests across participants, which corresponds to testing the cross-correlation at zero lag. Multiple comparisons across the 114 ROIs were controlled using the false discovery rate (FDR) at *p* < 0.05.

To further examine temporal dependencies beyond the canonical HRF assumption, we also repeated the cross-correlation and zero-lag *t*-tests using the *raw* (unconvolved) time series of pupil size and theory-of-mind (ToM) ratings. Finally, we performed a simple cross-correlation analysis between frame luminance and the group-level median pupil size time series for *The Present* video clip to explore direct visual–physiological coupling.

### 2.10. Finite Impulse Response Deconvolution

We performed FIR deconvolution to estimate the system response function between an input signal (*x(t)*) and an output signal (*y(t)*). Specifically, we first examined the pupillary system response using frame luminance as the input (*x(t)*) and pupil size as the output (*y(t)*).

Subsequently, we applied the same approach to estimate the HRFs in different brain regions using the various video-related features described above as input signals and fMRI time series from each ROI as outputs.

In general, the output signal can be modeled as the convolution of the input signal with an unknown system response function:

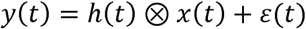

where (*h(t)*) denotes the system’s impulse response function, ⊗ represents the convolution operator, and (*ε(t)*) is the residual noise term.

The goal of FIR deconvolution is to estimate (*h(t)*) directly from the observed time series. We modeled (*h(t)*) as a finite-duration impulse response with a window of 20 seconds. Depending on the TR of the respective datasets, this window corresponded to a time-lagged design matrix containing 25 discrete lag regressors for The Present and Despicable Me (TR = 0.8 s), and 10 lag regressors for Partly Cloudy (TR = 2.0 s). The deconvolution was implemented using linear regression, where the input signal was expanded into a time-lagged design matrix containing shifted versions of (*x(t)*) at each time lag. The estimated regression coefficients correspond to the discrete samples of (*h(t)*).

To improve numerical stability and reduce overfitting, we first applied an autoregressive AR(1) pre-whitening step, followed by second-difference regularization with a penalty term (λ = 10). To ensure that our estimated response profiles were not overly smoothed or driven by this specific parameter choice, we conducted a sensitivity analysis by systematically testing a range of penalty values (λ = 0.1, 1, and 10). The criteria for robustness included the preservation of the estimated response shapes, the stability of the extracted peak latencies, and the spatial consistency of the group-level significance maps. Results across this tested range remained highly consistent, confirming the reliability of our chosen regularization parameter.

For the pupil–luminance analysis, FIR deconvolution was performed on the median pupil size time series. Both the pupil and luminance signals were z-scored before analysis to remove baseline offsets and normalize amplitudes.

For the fMRI data, FIR analysis was conducted separately for each participant and ROI. Prior to deconvolution, input features were resampled to match the fMRI sampling rate, and both input and output signals were z-scored.

### 2.11. Statistical analysis of FIR

To determine the statistical significance of the estimated FIR models, we utilized an *F*-test at the individual participant level to evaluate the overall variance explained by the full set of time-lagged predictors. To perform group-level statistical inference, we applied Fisher’s method to aggregate the individual-level *p*-values across all participants for each ROI (Fisher, 1925). The resulting group-level *p*-values were then corrected for multiple comparisons across the 114 ROIs using the FDR at *p* < 0.05, allowing us to systematically identify regions with significant FIR effects.

For regions demonstrating significant group-level responses, we obtained group-level response functions by averaging the individual FIR estimates after temporal alignment. To accurately quantify temporal hierarchies, we characterized the temporal dynamics by extracting the delay of the peak response (time-to-peak) for each ROI. To ensure that this metric captured the true primary physiological response while robustly resolving complex multiphasic profiles—such as initial dips or prominent undershoots—we evaluated the cumulative signal energy of the estimated FIR kernel. Specifically, the response function was bounded within a biologically plausible window, smoothed using a gentle Gaussian filter, and its positive and negative signal energies were computed by summing the squared amplitudes of each respective phase. The phase exhibiting the higher total energy dictated the dominant response polarity (positive activation or negative deactivation). The peak delay was then identified by extracting the time-to-peak of the global maximum for dominantly positive responses, or the global minimum for dominantly negative responses. These resulting temporal profiles, peak polarities, and peak latencies were then evaluated to compare physiological and neural dynamics across brain systems.

To evaluate the group-level reliability and stability of the estimated response shapes, we conducted a split-half consistency analysis. For each stimulus feature and movie condition, participants were first sorted by age and then assigned to two independent cohorts based on even and odd sample indices to ensure balanced age distributions between the halves. Within each split-half cohort, individual FIR estimates were averaged across participants to generate two independent group-level response profiles. The temporal consistency of the response shape was then quantified by calculating the Pearson correlation coefficient (r) between these two split-half time series for each ROI. This approach allowed us to map the spatial distribution of universally stereotyped versus highly idiosyncratic temporal profiles across the brain.

Finally, to complement this shape-based reliability metric and assess individual-level response uniformity, we performed a cross-individual sign consistency analysis. For each ROI, we evaluated the overall response polarity (positive versus negative deviation from baseline) of the estimated FIR function for each individual participant relative to the dominant group-level sign. Sign consistency for a given ROI was quantified as the proportion of participants whose individual response polarity matched the dominant group-level sign at that specific lag. This metric evaluates the directionality of the BOLD fluctuations across the sample, where a value of 0.5 represents baseline chance uniformity and higher values reflect a tightly synchronized, directional response across individuals.

## 3. Results

### 3.1. Sensory features: Temporal dynamics and spatial selectivity

Across all three movies, cross-correlation profiles exhibited a wide temporal spread across multiple lags (Supplementary Figure S1), likely reflecting the high inherent autocorrelation of continuous naturalistic stimuli and the sluggish BOLD signal. Despite this temporal smearing, the strongest zero-lag alignments reflected classical functional specialization, localizing primarily to modality-specific sensory cortices.

However, extended lag profiles revealed distinct temporal architectures depending on the complexity of the sensory feature. While basic features like frame luminance exhibited restricted spatial and temporal profiles mainly in visual areas (Figure 3A), more complex features—specifically frame contrast and auditory pitch—demonstrated organized temporal delay structures propagating into broader cortical networks (Figure 4A). These systematic temporal offsets across non-primary associative regions suggest that fixed, uniform HRF models may not fully capture the sequential dynamics of complex sensory processing.

**Figure 3.**
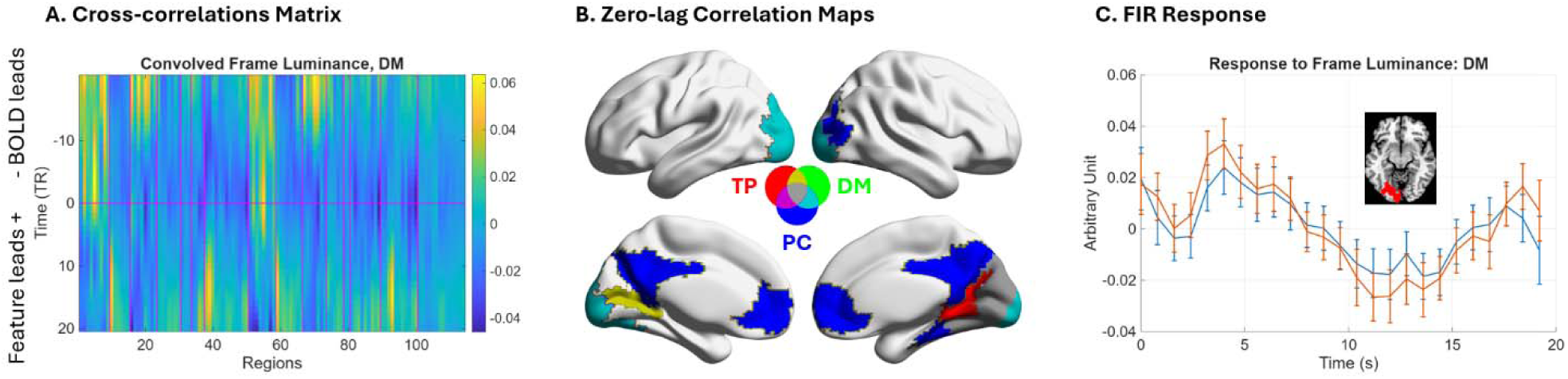
Neural correlates of frame luminance and temporal response dynamics. **(A)** Cross-correlation matrix illustrating the relationship between canonical HRF-convolved frame luminance and fMRI time series across 114 regions of interest (ROIs) for the movie *Despicable Me* (DM), plotted across multiple temporal lags. A positive lag indicates that the stimulus/feature time series precedes the BOLD signal (i.e., Feature lead), whereas a negative lag indicates that the BOLD signal precedes the feature (i.e., BOLD lead). **(B)** Cortical maps displaying significant positive zero-lag correlations between canonical HRF-convolved frame luminance (mean pixel intensity) and fMRI activity for *The Present* (TP), DM, and *Partly Cloudy* (PC). Statistical maps in panel B are thresholded at a false-discovery rate (FDR) of *p* < 0.05. **(C)** Estimated FIR response functions for ROIs exhibiting significant effects in the movie DM. The inset highlights the characteristic hemodynamic profiles of two primary visual regions. Error bar represents 95% confidence interval.

**Figure 4.**
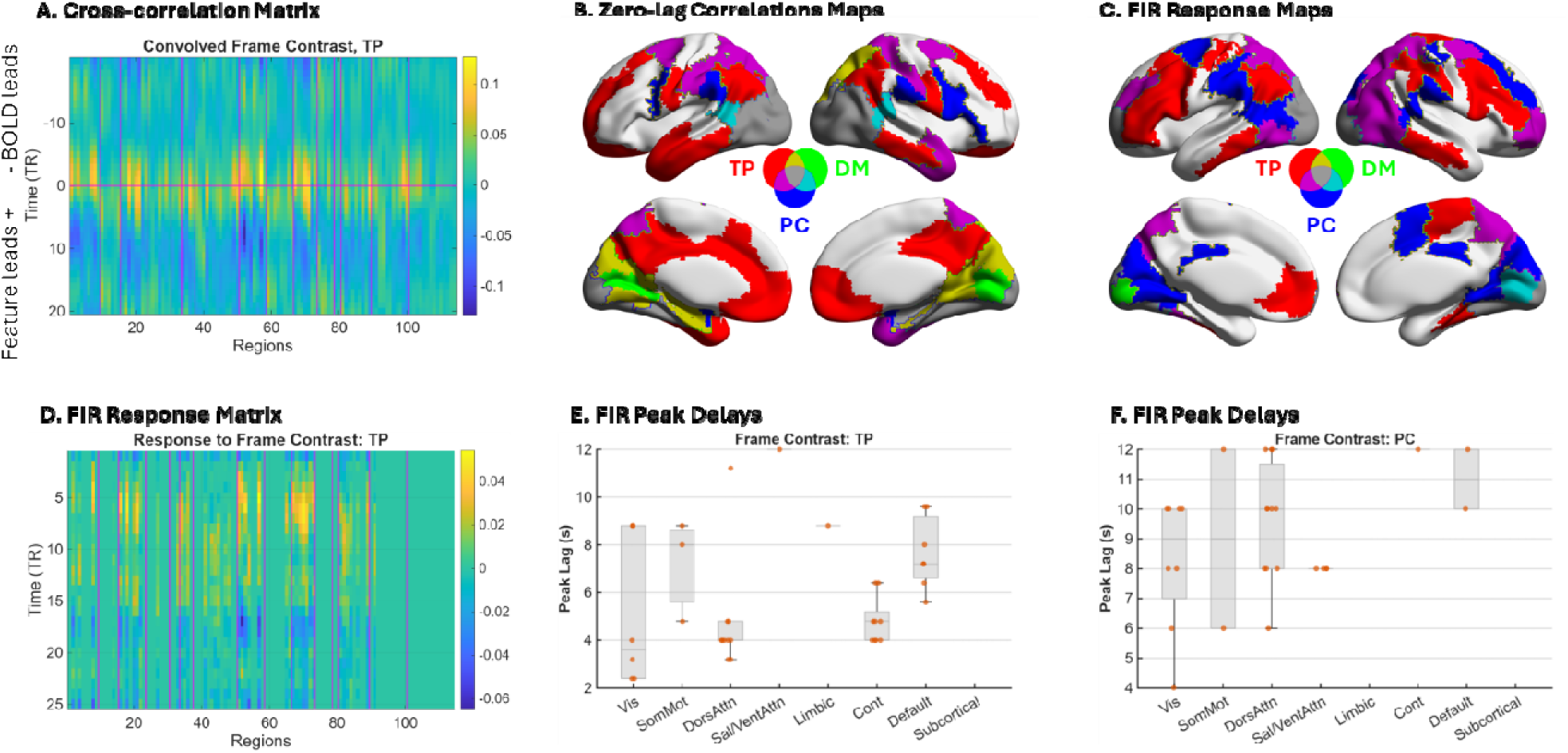
Neural correlates of frame contrast and associated temporal response dynamics. **(A)** Cross-correlation matrix illustrating the relationship between canonical HRF-convolved frame contrast and fMRI time series across 114 regions of interest (ROIs) for the movie *The Present* (TP) across multiple temporal lags. A positive lag indicates that the stimulus/feature time series precedes the BOLD signal (i.e., Feature lead), whereas a negative lag indicates that the BOLD signal precedes the feature (i.e., BOLD lead). **(B)** Cortical maps displaying significant positive zero-lag correlations between canonical HRF-convolved frame contrast and fMRI activity for TP, *Despicable Me* (DM), and *Partly Cloudy* (PC). **(C)** Cortical maps illustrating significant regional responses to frame contrast estimated via Finite Impulse Response (FIR) deconvolution. **(D)** Estimated FIR response functions across all 114 ROIs during movie TP, reflecting the diversity of hemodynamic profiles. **(E, F)** Distributions of estimated temporal delays (time-to-peak) for regions exhibiting positive FIR responses during movie TP (E) and PC (F), categorized across Yeo’s seven functional networks. All statistical maps (Panels B and C) are thresholded at a false-discovery rate (FDR) of p < 0.05.

#### 3.1.1. Luminance processing aligns with canonical assumptions

For frame luminance, prominent correlation peaks near zero lag suggested that the canonical HRF adequately approximates its hemodynamic delay. Consistent with prior literature, robust zero-lag correlations localized to primary and secondary visual cortices across all three movies, confirming the expected coupling between luminance fluctuations and early visual processing (Figure 3B).

FIR deconvolution yielded a more restricted set of significant regions compared to the zero-lag analysis (Supplementary Figure S2), likely reflecting the reduced statistical power of the unconstrained FIR model relative to a single-regressor approach. Nevertheless, significant FIR effects within the visual cortex during *Despicable Me* produced an estimated response function closely resembling the canonical hemodynamic profile (Figure 3C). Collectively, these results indicate that for basic sensory features, the standard HRF model remains a robust and sensitive methodological approach.

#### 3.1.2. Visual contrast and the complexity of naturalistic narrative

Zero-lag correlations for frame contrast elicited a predominantly modality-specific spatial profile, localized primarily within visual cortices across the stimuli (Figure 4B). However, cross-correlation profiling across extended lags demonstrated that naturalistic visual contrast effects extend well beyound isolated sensory regions. Specifically, the cross-correlation matrix for *The Present* (TP) revealed a widespread footprint extending well beyond primary sensory areas (Figure 4A), suggesting the sequential engagement of higher-order functional networks.

FIR deconvolution further mapped this widespread temporal engagement (Figure 4C; Supplementary Figure S3). While the spatial overlap of significant FIR effects across all three movies was constrained to primary visual regions (largely due to a more limited spatial response during *Despicable Me*), the unconstrained FIR model successfully captured a distinct temporal architecture across the broader cortex for both *The Present* and *Partly Cloudy*.

The estimated FIR response functions exhibited a clear, organized temporal progression across varying ROIs (Figure 4D). When quantifying this temporal hierarchy by extracting positive peak response latencies across Yeo’s seven functional networks (Figure 4E, 4F), both movies demonstrated a systematic delay structure. Visual networks consistently exhibited the shortest response latencies, whereas progressive delays were observed in higher-order networks, culminating in the longest latencies within the somatomotor, dorsal attention, ventral attention/salience, and default mode networks.

#### 3.1.3. Auditory pitch and stimulus collinearity

Similar to visual features, zero-lag correlations for auditory pitch elicited strong, modality-specific alignments within core auditory cortices. However, during *Despicable Me*, these significant correlations extended substantially into broader cortical networks (Figure 5A).

**Figure 5.**
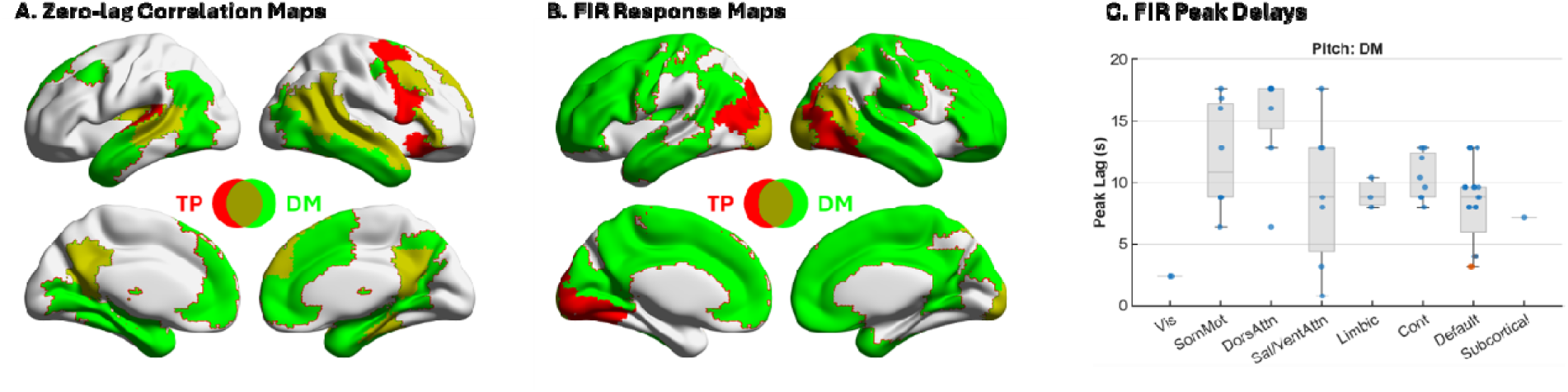
Neural correlates of auditory pitch and temporal delay distributions. **(A)** Cortical maps displaying positive zero-lag correlations between the canonical HRF-convolved auditory pitch and fMRI activity during *The Present* (TP) and *Despicable Me* (DM). **(B)** Cortical maps illustrating significant regional responses to auditory pitch estimated via Finite Impulse Response (FIR) deconvolution for both videos. Statistical maps in panels A and B are thresholded at a false-discovery rate (FDR) of p < 0.05. **(C)** Distributions of estimated temporal delays (time-to-peak) for regions exhibiting positive FIR responses, categorized across Yeo’s seven functional networks. Red markers denote two core auditory regions that are assigned to the Default Mode Network in the Yeo parcellation.

FIR deconvolution revealed complex, stimulus-dependent temporal response profiles (Supplementary Figure S4). During *The Present*, regions exhibiting significant FIR effects were unexpectedly localized predominantly to visual cortices (Figure 5B). Extracted response functions in these visual areas revealed a distinct negative signal deflection within the first 5 seconds post-stimulus. This initial negative temporal profile indicates that auditory pitch likely covaries inversely with concurrent salient visual features specific to this stimulus, highlighting the bundled nature of cross-modal naturalistic stimulus rather than direct auditory processing in the visual cortex.

Conversely, FIR deconvolution for *Despicable Me* identified widespread significant cortical effects distributed across multiple functional networks (Figure 5B). Mirroring the temporal architecture observed for frame contrast, the latencies of positive peak responses to auditory pitch exhibited a hierarchical progression. The shortest latencies localized to core auditory regions, followed by sequentially increasing delays systematically progressing through the dorsal attention, somatomotor, ventral attention/salience, limbic, executive control, and ultimately the default mode networks. (Note: Core auditory cortices are assigned to the default mode network boundary in the standard Yeo parcellation based on resting-state data; to maintain anatomical precision, these specific nodes are denoted with red markers in Figure 5C).

### 3.2. Pupil size: Biological delays and autonomic network coupling

Pupil size dynamics exhibited robust inter-individual consistency across the sample (mean ISC r = 0.63). As expected, the group-median pupil trace demonstrated a strong inverse relationship with frame luminance, with pupil constriction lagging luminance changes by approximately 0.67 seconds (Figure 6A and Supplementary Figure S5).

**Figure 6.**
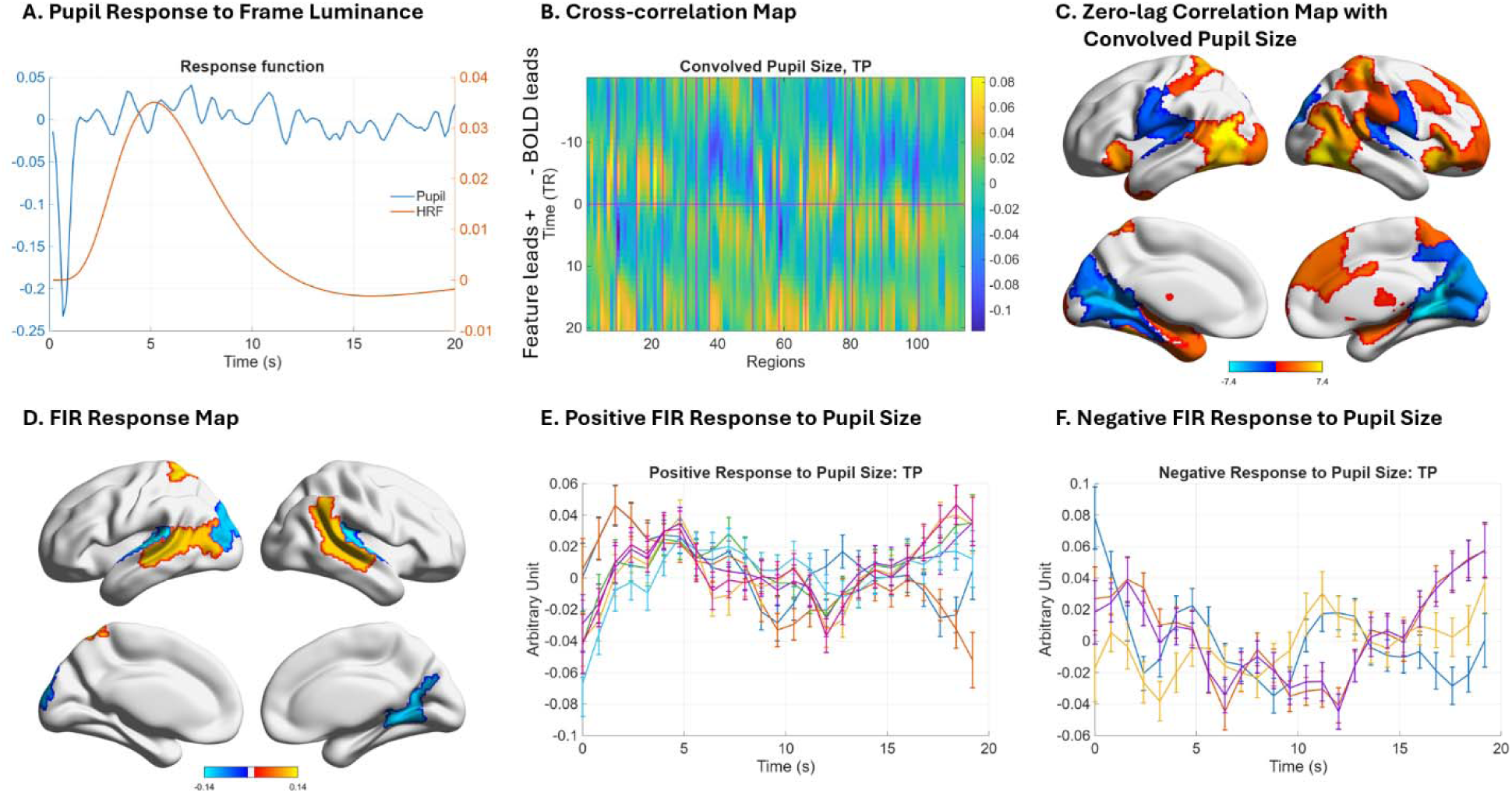
Neural correlates of pupil size and associated temporal response dynamics for movie *The Present* (TP). **(A)** Temporal response function of pupil size relative to frame luminance, shown in comparison to a canonical hemodynamic response function (HRF). **(B)** Cross-correlation matrix illustrating the relationship between the raw pupil size and fMRI time series across 114 regions of interest (ROIs) for movie TP across multiple temporal lags. **(C)** Cortical maps displaying significant positive (hot) and negative (cold) zero-lag correlations between canonical HRF-convolved pupil size and fMRI activity for movie TP. A positive lag indicates that the stimulus/feature time series precedes the BOLD signal (i.e., Feature lead), whereas a negative lag indicates that the BOLD signal precedes the feature (i.e., BOLD lead). **(D)** Cortical maps illustrating significant regional positive (hot) and negative (cold) responses to pupil size estimated via Finite Impulse Response (FIR) deconvolution. **(E) and (F)** Estimated FIR response functions for significant positive and negative response regions, respectively. All statistical maps (Panels C, D, and E) are thresholded at a false-discovery rate (FDR) of *p* < 0.05. Error bar represents 95% confidence interval.

Given this relatively short physiological delay, cross-correlation profiling using the standard HRF-convolved pupil trace revealed a slight temporal mismatch: the convolved trace lagged marginally behind regional BOLD responses (Figure 6B). This slight phase shift indicates a minor temporal overcompensation—a "double-delay"—introduced by applying the canonical HRF to a peripheral signal that already contains an inherent biological latency. Despite this slight shift, zero-lag correlations successfully captured widespread networks, revealing positive correlations in lateral occipitotemporal, lateral frontoparietal, subcortical, and insular regions, alongside negative correlations in medial visual and motor cortices (Figure 6C).

To capture these temporal dynamics without HRF overcompensation, we applied FIR deconvolution. While this unconstrained model yielded fewer significant regions overall, it successfully mapped response profiles predominantly within temporo-occipital regions (Figure 6D). Because the estimated FIR trajectories were highly complex and multiphasic (Supplementary Figure S6)—likely reflecting the bundled dynamics of autonomic arousal and highly correlated visual processing—we classified the dominant response polarity of each region using its cumulative signal energy. Regions exhibiting a dominantly positive response to pupil size included the bilateral temporal cortices (Figure 6E), whereas dominantly negative responses were localized to the bilateral peri-Sylvian and medial visual regions (Figure 6F).

### 3.3. Theory of Mind ratings: Context-dependent dynamics of social cognition

Inter-individual correlations of ToM rating time series demonstrated high consistency across *The Present* (*r* = 0.55), *Despicable Me* (*r* = 0.36), and *Partly Cloudy* (*r* = 0.56; all *p* < 0.001), validating the use of the median rating time series for subsequent fMRI analyses. When mapping BOLD correlates using the standard HRF-convolved ToM ratings, cross-correlation profiles revealed substantial temporal misalignments that varied heavily by stimulus (Supplementary Figure S7).

For *The Present*, peak correlations within the visual and dorsal attention networks occurred at positive lags (BOLD leading; Supplementary Figure S7A), indicating that canonical convolution may overcompensated for the inherent delays in the ToM ratings. For *Partly Cloudy*, cross-correlation peaks in several visual areas aligned closely with the zero-lag line (Supplementary Figure S7C). The results for *Despicable Me* were more complex (Supplementary Figure S7B); while visual regions lacked clear zero-lag peaks, auditory regions exhibited massive feature-leading delays of more than 10 seconds. Because the convolved time series already incorporates an ∼6-second hemodynamic delay, these extended lags cannot be explained by neurovascular coupling alone, and instead likely reflect the collinearity of ToM with slowly unfolding narrative features. Consequently, zero-lag correlation maps using the HRF-convolved ToM ratings revealed only one consistent significant cluster in the left temporo-occipital region across all three videos (Figure 7A).

**Figure 7.**
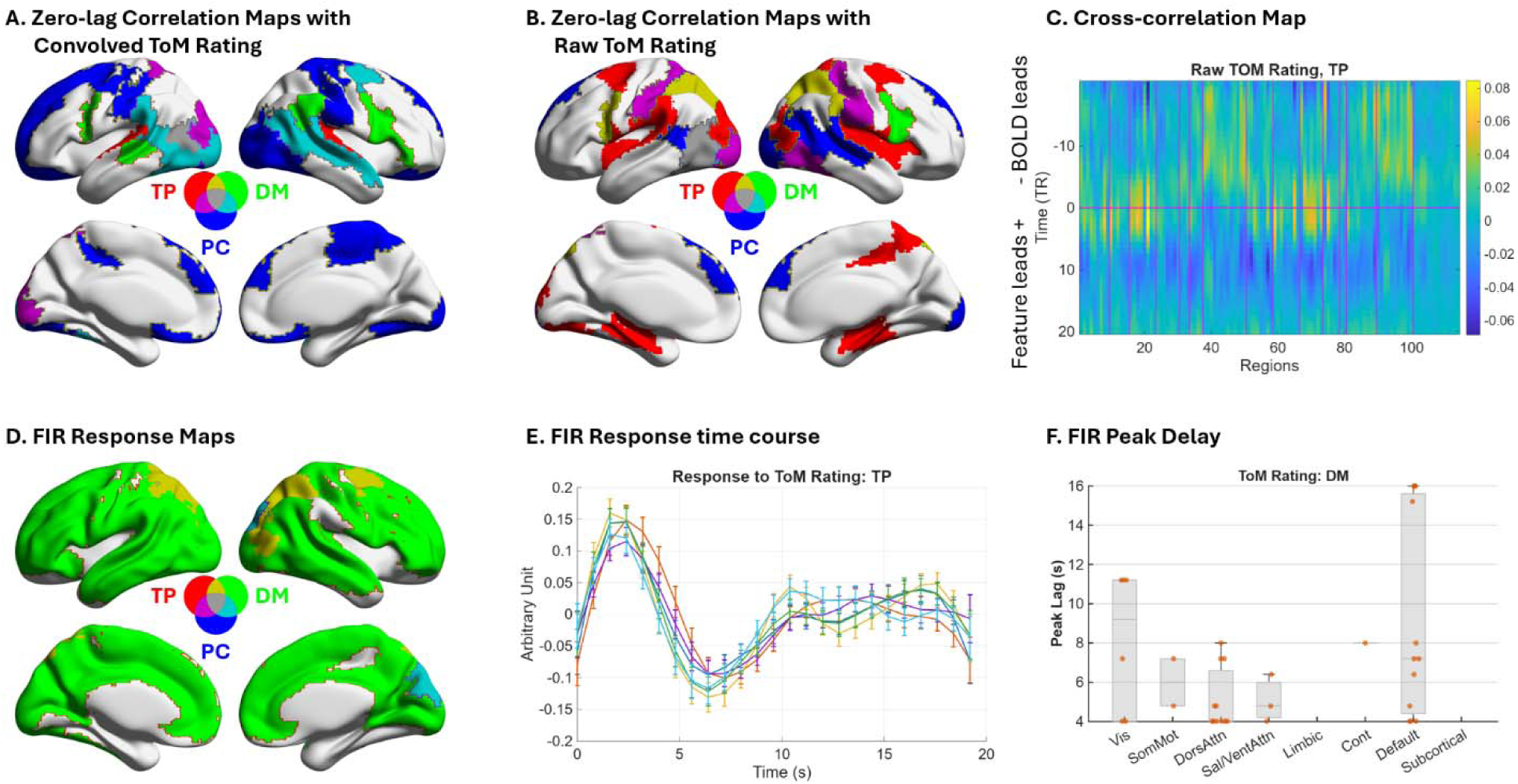
Neural correlates of Theory of Mind (ToM) ratings and associated temporal response dynamics. **(A)** Cortical maps displaying significant positive zero-lag correlations between canonical HRF-convolved ToM ratings and fMRI activity for *The Present* (TP), *Despicable Me* (DM), and *Partly Cloudy* (PC). **(B)** Cortical maps displaying significant positive zero-lag correlations between raw (unconvolved) ToM ratings and fMRI activity across the three videos. **(C)** Cross-correlation matrix illustrating the relationship between the raw ToM rating and fMRI time series across 114 regions of interest (ROIs) for movie TP across multiple temporal lags. A positive lag indicates that the stimulus/feature time series precedes the BOLD signal (i.e., Feature lead), whereas a negative lag indicates that the BOLD signal precedes the feature (i.e., BOLD lead). **(D)** Cortical maps illustrating significant regional responses to ToM ratings estimated via Finite Impulse Response (FIR) deconvolution. **(E)** Estimated FIR response functions for significant ROIs during movie TP, highlighting the rapid peak latencies. **(F)** Distributions of estimated temporal delays (time-to-peak) for regions exhibiting positive FIR responses during movie DM, categorized across Yeo’s seven functional networks. All statistical maps (Panels A, B, and D) are thresholded at a false-discovery rate (FDR) of *p* < 0.05. Error bar represents 95% confidence interval.

Because continuous behavioral ratings intrinsically incorporate cognitive processing and motor execution delays, applying the canonical HRF might artifactually "double-delay" and over-smooth the signal. To address this, we repeated the cross-correlation analyses using the raw, unconvolved ToM ratings. This approach yielded more temporally aligned cross-correlation profiles, with peaks shifting much closer to zero lag for both *The Present* (Figure 7C) and *Partly Cloudy* (Supplementary Figure S8F). Consequently, the zero-lag correlation maps using the raw ToM time series revealed a greater spatial overlap of significant clusters across the three videos, particularly in the left occipito-temporal junction (Figure 7B).

While using the raw ToM time series avoids the penalty of double-smoothing, it still rigidly assumes uniform neuro-behavioral latency across all brain regions and stimuli. FIR deconvolution revealed that ToM processing dynamics are stimulus-dependent (Figure 7D and Supplementary Figure S8). For *The Present*, significant FIR effects were restricted primarily to the dorsal visual stream and parietal regions. Crucially, the extracted response functions in these regions peaked extremely rapidly (1.6–2.4 seconds; Figure 7E), demonstrating empirically why the canonical HRF model (which assumes a ∼5-second peak) was systematically inappropriate for this specific stimulus.

Conversely, FIR deconvolution for *Despicable Me* revealed widespread significant cortical responses. Unlike the distinct temporal hierarchies observed during basic sensory processing, the estimated FIR delays for ToM in *Despicable Me* did not exhibit a clear cortical feed-forward pattern (Figure 7F). This widespread, complex temporal profile likely captures the bundled dynamics of mentalizing required for this specific narrative*-*reflecting the distributed, simultaneous integration of dialogue, abstract social rules, and multi-character dynamics.

Together, these results demonstrate that flexible approaches like FIR deconvolution are essential to capture the highly context-dependent speeds of social cognition during naturalistic viewing.

### 3.4. FIR consistency analysis

To further investigate the widespread cortical engagement observed in the initial FIR analyses, we mapped the spatial distribution of split-half temporal consistency (Supplementary Figure S9). Across most cortical networks, split-half consistency of the estimated response shapes was typically high (r > 0.6). Sensory regions, in particular, exhibited exceptionally strong split-half consistency during corresponding stimulus conditions—such as the visual cortex during frame contrast (r > 0.9)—pointing to a stereotyped temporal response across subjects.

In contrast, within conditions where significant FIR activations initially appeared widespread (such as auditory pitch or ToM during *Despicable Me*), higher-order associative areas—including the default mode network and lateral prefrontal regions—demonstrated a sharp drop in consistency, falling to r = 0.2 to 0.3 (Figure 8). This spatial divergence indicates that while the unconstrained FIR model detects significant non-zero variance in these associative networks, the actual shape of the estimated temporal response is highly variable across individuals. Consequently, these widespread frontal effects likely reflect idiosyncratic, subject-specific fluctuations or model overfitting rather than a uniform, canonical hemodynamic profile.

**Figure 8.**
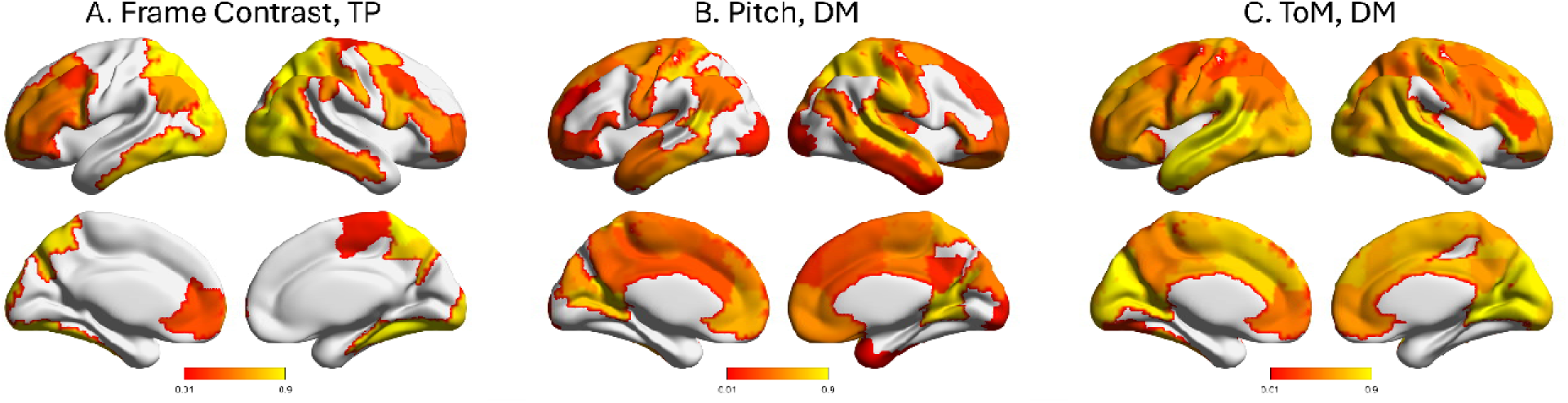
Cross-individual split-half consistency of finite impulse responses (FIR) for three representative conditions: frame contrast in TP (*The Present*), auditory pitch in DM (*Despicable Me*), and Theory of Mind (ToM) in DM.

Finally, we evaluated the absolute sign consistency (directionality) of the FIR estimates across individuals. This analysis revealed notably lower uniformity across participants relative to the split-half correlations (Supplementary Figure S10). This divergence indicates that while the relative timing and overall trajectory of the feature-evoked responses are generally conserved across at group level, the absolute baseline polarity and scaling of individual-level FIR estimates are highly vulnerable to idiosyncratic variation and subject-specific noise.

## 4. Discussion

In the present study, we investigated the temporal dynamics of BOLD responses to continuous naturalistic features during movie watching. By integrating multi-level features—low-level sensory properties, physiological (pupil size), and subjective cognitive annotations (ToM)—we evaluated how well conventional hemodynamic modeling captures the brain’s nested temporal hierarchies. Using cross-correlation and FIR deconvolution, we mapped the unconstrained BOLD response profiles across the whole brain.

Our main findings are threefold. First, while canonical HRF models adequately capture sensory feature dynamics, they systematically misalign physiological and cognitive signals. Second, convolving inherently delayed signals (e.g., pupil size) with a standard HRF introduces a redundant “double-delay,” causing a phase mismatch with the peak of neurovascular coupling. Finally, FIR deconvolution revealed complex, multiphasic regional response. Rather than reflecting isolated feature processing, these profiles capture the bundled dynamics of correlated naturalistic dimensions, highlighting a progression of processing delays from primary sensory cortices to higher-order associative networks. Together, these results underscore the necessity of flexible temporal modeling to accurately interpret the diverse processing timescales engaged during naturalistic viewing.

### 4.1. Sensory features

For low-level sensory features (frame luminance, contrast, and pitch), both cross-correlation and FIR analyses confirmed that the canonical HRF effectively captured the temporal delay between stimulus dynamics and BOLD responses. Peak correlations consistently occurred at lags of 4–6 seconds, and FIR-estimated impulse response in modality specific brain regions mirrored the canonical HRF’s characteristic rise, peak, and undershoot phases. This aligns with both classic event-related and naturalistic fMRI studies (Boynton et al., 1996; Glover, 1999), confirming that neurovascular coupling in early sensory regions is highly stereotyped and well-approximated by standard models.

However, our unconstrained FIR models also revealed systematic inter-regional delays propagating from primary sensory areas to higher-order associative and default mode networks. This temporal progression provide empirical support for the theoretical framework of hierarchical cortical timescales, or TRWs (Lerner et al., 2011; Meer et al., 2020). Early sensory cortices possess short TRWs adapted for rapid physical changes, yielding sharp FIR profiles and early peak latencies. In contrast, higher-order networks-tasked with integrating sensory input into semantic and narrative context-exhibit longer TRWs, reflected in significantly slower, temporally diffuse BOLD responses (Lewis et al., 2018). Applying a uniform canonical HRF across the entire brain obscures this fundamental hierarchical topography, underscoring the value of flexible temporal modeling to capture varying cortical integration timescales.

### 4.2. Pupil size

Previous fMRI studies have often modeled pupil diameter as a physiological regressor by convolving it with a canonical HRF before correlating it with BOLD signals (Schneider et al., 2016; Yellin et al., 2015). This approach, however, conflates two distinct system response functions: the HRF, which characterizes local neurovascular coupling, and the Physiological Response Function (PRF), which governs the upstream somatic dynamics transforming external stimuli or internal autonomic states into peripheral physiological measures. Convolving raw pupil size with a canonical HRF implicitly assumes it acts as an instantaneous neural proxy. However, our analysis revealed that pupil constriction intrinsically lags luminance changes by 0.67 to 1.0 seconds, reflecting the established biological latency of the pupillary and autonomic responses (Mathot, 2018).

While this relatively short physiological delay permits standard HRF-convolved models to successfully map widespread activation networks (Figure 6C), our cross-correlation profiles revealed a systematic temporal mismatch. Because the pupil trace already contains a ∼1-second PRF delay relative to the initial stimulus-driven neural event, applying a full canonical HRF convolution overcompensates, introducing a minor "double-delay." Consequently, standard GLM approaches utilizing the canonical HRF miss the absolute peak of this temporal coupling.

By employing FIR deconvolution, we captured these temporal dynamics without HRF overcompensation. Rather than reflecting an isolated pupillary mechanism, the extracted multiphasic response profiles capture the bundled dynamics of autonomic arousal and high correlated visual processing. We demonstrated that bilateral temporal cortices exhibit a dominantly positive response to pupil size, whereas medial visual and motor regions exhibit dominantly negative responses.

From a neurophysiological perspective, pupil diameter serves as a peripheral marker of central arousal, heavily influenced by the locus coeruleus–norepinephrine (LC–NE) system (Aston-Jones and Cohen, 2005; Joshi et al., 2016; Murphy et al., 2014). The widespread correlations observed between raw pupil fluctuations and BOLD activity in the insula, temporoparietal junction, and thalamus align with the distributed targets of the LC–NE network (Breton-Provencher and Sur, 2019; Shine et al., 2019). Thus, these unconstrained response profiles likely reflect the integrated, total effect of neuromodulatory arousal and adaptive cognitive control during naturalistic viewing.

### 4.3. Theory of Mind Rating

In this study, an independent cohort provided continuous ratings reflecting their perceived understanding of the characters’ mental states, intended to index moment-to-moment engagement of ToM (Frith and Frith, 2006; Schurz et al., 2021). Unlike low-level sensory or physiological features, the temporal coupling between such introspective, socially grounded judgments and underlying neurovascular activity is far less constrained and varies heavily with narrative context (Jacoby et al., 2016; Richardson et al., 2018).

Generating a continuous introspective ratings inherently integrate information over several seconds due to cognitive evaluation and motor execution delays (Nummenmaa et al., 2012; Zahn et al., 2009). Because this behavioral trace is intrinsically a temporally smoothed and delayed process, applying an additional canonical HRF convolution is mathematically redundant. This secondary convolution compounds the temporal dynamics, creating a "double-delay" that fundamentally shifts the regressor out of phase with the unconstrained neural activity while over-smearing the signal variance. Consequently, our analysis demonstrated that using the raw, unconvolved ToM ratings yielded significantly improved spatial consistency across the different movie clips compared to the HRF-convolved approach.

Furthermore, performing FIR deconvolution directly on this raw trace logically yields an accelerated estimated hemodynamic peak, as the behavioral latency already accounts for a portion of the typical neurovascular lag. This empirically demonstrates why the canonical HRF model, which rigidly assumes a ∼5-second peak, is systematically inappropriate when modeling inherently delayed behavioral traces.

Finally, our FIR results revealed that ToM processing dynamics are context-dependent. While sensory features exhibited a clear sequential propagation from primary to higher-order networks, ToM processing during complex narratives (e.g., Despicable Me) did not. Instead, the estimated temporal delays were widespread and synchronous across functional networks. Rather than reflecting an isolated, sequentially hierarchical ToM pathway, this distributed profile likely captures the bundled effects of concurrent narrative integration—reflecting the simultaneous processing of dialogue, abstract social rules, and dynamic multi-character interactions.

### 4.4. Complexity of naturalistic stimuli

A striking observation in our analysis was the substantial variability in brain responses across the three films. While sensory features generally exhibited consistent activations, high-level cognitive features like ToM did not uniformly engage classical mentalizing networks across all datasets. This divergence likely stems from differences in narrative complexity: *Despicable Me* involves complex, dialogue-driven social rules (yielding widespread cortical engagement), whereas short animations like *The Present* or *Partly Cloudy* rely on non-verbal visual cues (engage a restricted social-perceptual sub-network). This inter-stimulus variability is further compounded by differences in data acquisition, such as varying temporal resolutions and data lengths. Given these discrepancies, cross-dataset comparisons are inherently exploratory. However, by leveraging this heterogeneity, we successfully identified canonical response profiles that persist across diverse naturalistic viewing conditions.

A final critical methodological consideration when mapping these temporal dynamics is the inherent collinearity of naturalistic stimulus features. Unlike highly controlled experiments, cinematic features are continuously presented and frequently covary with unmeasured features (e.g., semantic meaning or emotional valence). Because we largely evaluated the features one at a time, our unconstrained FIR profiles reflect the total effect of bundled stimulus dimensions rather than isolated feature processing. For instance, the widespread cortical responses to auditory pitch likely capture its natural covariance with human speech and prosody. While combining multiple features into a single, comprehensive general linear model could theoretically isolate the unique variance, expanding numerous correlated features into dozens of time-lagged predictors severely inflates variance and destabilizes FIR estimates. Future studies employing advanced regularization techniques (e.g., ridge regression) or dimensionality reduction are necessary to disentangle these overlapping profiles.

Beyond regularized frameworks, a compelling alternative to handling stimulus complexity is the use of purely data-driven discovery methods. Techniques such as independent component analysis (ICA), tensor component analysis (TCA), Hidden Markov Models (HMMs; (Baldassano et al., 2017)), response time series clustering, and principal component analysis (PCA) (Di and Biswal, 2022) bypass the need for predefined feature annotations or rigid hemodynamic assumptions. These unsupervised approaches excel at isolating intrinsic, latent brain states directly from the unconstrained BOLD data. However, mapping those extracted components back to specific narrative events often requires post-hoc interpretation. Thus, our feature-driven approach and blind data-driven decompositions provide complementary windows into naturalistic brain function.

### 4.5. Methodological limitations

A key methodological limitation of this study is the integration of proxy data across partially overlapping participant cohorts. Correlating physiological (pupil size) and behavioral (ToM) time series from independent groups with fMRI BOLD signals relies on the assumption that stimulus-driven cognitive and autonomic dynamics are highly homogeneous across individuals. While our split-half consistency metrics justify this group-level proxy, pairing data across distinct participant sets inherently introduces unmeasured between-subject variance into the model error term. Consequently, our models estimate generalized, stimulus-driven group responses rather than true within-individual cognitive-neural coupling. Future investigations incorporating simultaneous in-scanner tracking are essential for disentangling shared, stimulus-driven responses from subject-specific neurovascular variations.

Furthermore, while a notable advantage of FIR deconvolution is its ability to detect atypical delayed responses, its unconstrained nature carries a high risk of overfitting. For certain low-level features (e.g., frame contrast), the FIR omnibus tests suggested near whole-brain engagement. By mapping the spatial distribution of split-half consistency, we were able to disambiguate two distinct drivers of these widespread effects. In primary sensory cortices, high temporal consistency confirmed that the responses were both significant and uniformly stereotyped across the population. Conversely, in widespread frontal and associative regions, temporal consistency dropped markedly. This suggests that the FIR model, possessing multiple degrees of freedom, can capture highly idiosyncratic, subject-specific fluctuations or overfit structured noise in these higher-order areas. Future naturalistic fMRI studies utilizing unconstrained temporal modeling must report response reliability alongside standard activation maps to accurately differentiate true, shared stimulus processing from model overfitting.

### 4.6. Broader implications for hemodynamic response

Traditional HRF estimation approaches have primarily relied on event-related or block designs (Dale and Buckner, 1997; Glover, 1999). While instrumental, they artificially constrain the complexity of real-world experiences. In contrast, our analysis provides robust empirical evidence that hemodynamic responses can be estimated directly from continuous naturalistic data, encompassing sustained attention, emotional engagement, and social inference (Hasson et al., 2004; Nastase et al., 2019). The flexible FIR model revealed systematically diverse temporal profiles across regions, consistent with prior evidence that HRF dynamics vary with vascular and neuronal properties (Aguirre et al., 1998; Handwerker et al., 2004; Lindquist et al., 2009).

Because this flexibility comes at the cost of increased variance (Goutte et al., 2000; Pedregosa et al., 2015), an important priority for future research is to assess the inter-individual variability of HRF estimates derived from continuous data. Individual differences in baseline physiology may systematically influence HRF latency and shape (Handwerker et al., 2004; West et al., 2019). Quantifying these sources of variability could enhance reproducibility and between-subject interpretability, with potential applications in aging (Chen et al., 2024; West et al., 2019) and cerebrovascular disease.

Finally, integrating FIR-based estimation with the analyses of multi-level narrative features illustrates the value of temporal modeling. The observation that sensory features align well with the canonical HRF, whereas physiological and introspective signals exhibit complex, delayed relationships, confirms that hemodynamic coupling differs systematically across temporal receptive windows. Data-driven HRF estimation provides both a methodological advance and a conceptual framework for understanding how distinct neural systems integrate information over time.

## 5. Conclusion

In summary, this study systematically characterized the temporal relationships between continuous sensory, physiological, and behavioral features and fMRI BOLD signals during naturalistic movie viewing. Using cross-correlation and unconstrained FIR deconvolution, we demonstrated that while low-level sensory features generally align with canonical hemodynamic expectations, slower physiological signals (pupil size) and high-level cognitive annotations (ToM) systematically deviate from these standard models. Crucially, applying a uniform canonical HRF to inherently delayed physiological and behavioral time series introduces redundant temporal shifts. Because signals like pupil size are governed by their own PRFs, standard HRF convolution overcompensates for their intrinsic biological latency, resulting in a "double-delay" that misaligns with the absolute peak of neurovascular coupling.

By employing data-driven FIR deconvolution, we successfully mapped unconstrained temporal response profiles and sequential processing hierarchies across the brain. Recognizing the inherent collinearity of naturalistic stimuli, these estimated profiles capture the bundled, multi-dimensional dynamics of real-world processing rather than perfectly isolated feature effects. Furthermore, by integrating a split-half consistency analysis, we provided a vital framework for differentiating universally stereotyped neural responses from idiosyncratic variations and model overfitting in higher-order networks. These findings highlight the necessity of flexible, reliability-tested temporal modeling when mapping the complex, multi-level processing timescales engaged during naturalistic paradigms.

## Acknowledgments

During the preparation of this revised manuscript, the authors utilized Gemini (Google) solely for language polishing and assistance with drafting technical summaries. No artificial intelligence tools were used in the generation of research data, data analysis, or the scientific interpretation of the results. The authors thoroughly reviewed and edited all AI-assisted text and take full responsibility for the final content of the manuscript.

## Data and Code Availability

MRI and pupil size data were obtained from publicly available sources: the Healthy Brain Network (HBN) dataset (http://fcon_1000.projects.nitrc.org/indi/cmi_healthy_brain_network/) and the OpenNeuro repository (https://openneuro.org/datasets/ds000228).

## Author Contributions

X.D. conceived and designed the study, developed the methodology, performed the analyses, and wrote the original draft of the manuscript. G.B.H. collected and curated the data and contributed to manuscript review and editing. B.B.B. and X.D. provided resources and secured funding for the project. All authors contributed to the interpretation of results and approved the final version of the manuscript.

