## Supplementary materials for "Beyond the Canonical HRF: Flexible Temporal Modeling Reveals Unconstrained BOLD Profiles During Naturalistic Viewing"

**Summary**

This supplementary document provides extended visualizations and statistical maps supporting the primary findings on the spatiotemporal dynamics of BOLD responses during naturalistic movie viewing. It includes comprehensive cross-correlation matrices and unconstrained Finite Impulse Response (FIR) deconvolution profiles for low-level sensory features (frame luminance, frame contrast, and auditory pitch), a physiological proxy of autonomic arousal (pupil size), and high-level cognitive annotations (Theory of Mind ratings). In addition to detailing unthresholded temporal profiles and group-level statistical maps, this document includes comprehensive cross-cohort reliability analyses (split-half and sign consistency). Together, these figures further demonstrate the widespread variation in hemodynamic delays across functional networks, highlight the complex, bundled nature of naturalistic stimulus processing, and emphasize the necessity of reliability-tested, flexible temporal modeling.

**Table of Contents**

- **Supplementary Figure S1.** Spatiotemporal cross-correlation matrices for sensory features
- **Supplementary Figure S2.** FIR deconvolution profiles for frame luminance
- **Supplementary Figure S3.** FIR deconvolution profiles for frame contrast
- **Supplementary Figure S4.** FIR deconvolution profiles for auditory pitch

- **Supplementary Figure S5.** Temporal relationship between frame luminance and pupillary dynamics
- **Supplementary Figure S6.** FIR deconvolution profiles for pupil size
- **Supplementary Figure S7.** Spatiotemporal cross-correlation matrices for convolved and raw Theory of Mind (ToM) ratings
- **Supplementary Figure S8.** FIR deconvolution profiles for theory of mind (ToM) rating
- **Supplementary Figure S9.** Cross-individual split-half consistency of finite impulse responses (FIR)
- **Supplementary Figure S10.** Cross-individual sign consistency of finite impulse responses (FIR)

**Supplementary Figure S1. Spatiotemporal cross-correlation matrices for sensory features.** Matrices illustrate the relationship between canonical HRF-convolved sensory features and fMRI activity across 114 regions of interest (ROIs) for the movies *The Present* (TP, first column), *Despicable Me* (DM, second column), and *Partly Cloudy* (PC, third column). Rows represent frame luminance (mean pixel intensity; top), frame contrast (pixel standard deviation; middle), and auditory pitch (bottom). A positive lag indicates that the stimulus/feature time series precedes the BOLD signal (i.e., Feature lead), whereas a negative lag indicates that the BOLD signal precedes the feature (i.e., BOLD lead). Vertical dividers delineate ROI assignments across the left and right hemispheres according to Yeo's seven-network parcellation, as well as subcortical regions. Each matrix reflects the extended temporal "footprint" of correlations across multiple lags.

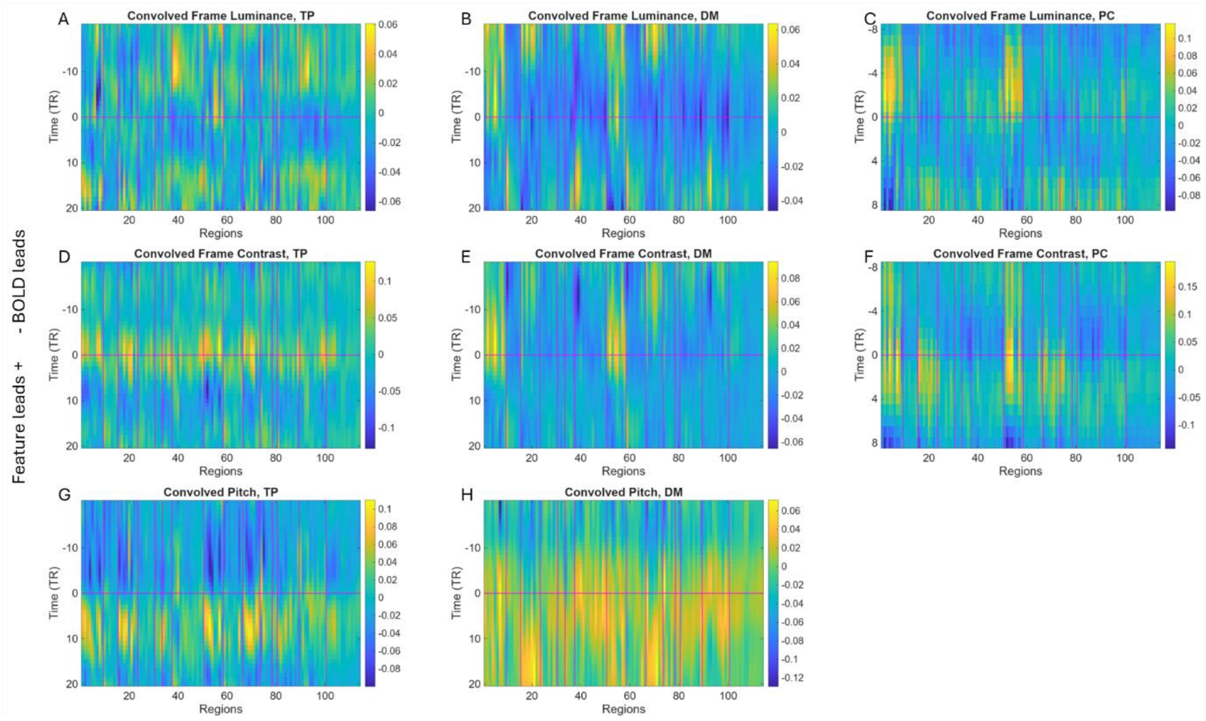

**Supplementary Figure S2. FIR deconvolution profiles for frame luminance.** Panels illustrate the estimated system response functions across 114 regions of interest (ROIs) for the movies *The Present* (TP, first column), *Despicable Me* (DM, second column), and *Partly Cloudy* (PC, third column). **Top row:** Unthresholded mean FIR response profiles across all 114 ROIs. **Bottom row:** Significant regional response functions identified through group-level statistical inference. Statistical significance was determined by aggregating individual-level $F$-tests using Fisher's method, with the resulting group-level maps thresholded at a false discovery rate (FDR) of $p < 0.05$.

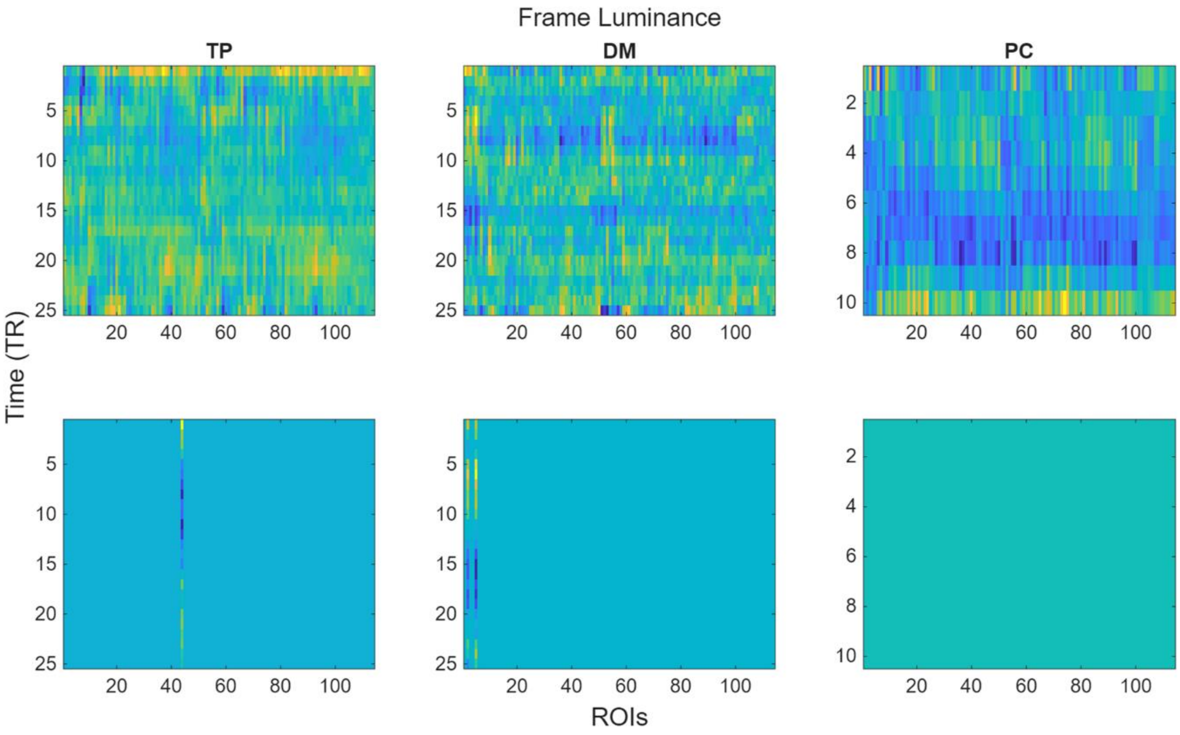

**Supplementary Figure S3. FIR deconvolution profiles for frame contrast.** Panels illustrate the estimated system response functions across 114 regions of interest (ROIs) for the movies *The Present* (TP, first column), *Despicable Me* (DM, second column), and *Partly Cloudy* (PC, third column). **Top row:** Unthresholded mean FIR response profiles across all 114 ROIs. **Bottom row:** Significant regional response functions identified through group-level statistical inference. Statistical significance was determined by aggregating individual-level *F*-tests using Fisher's method, with the resulting group-level maps thresholded at a false discovery rate (FDR) of $p < 0.05$.

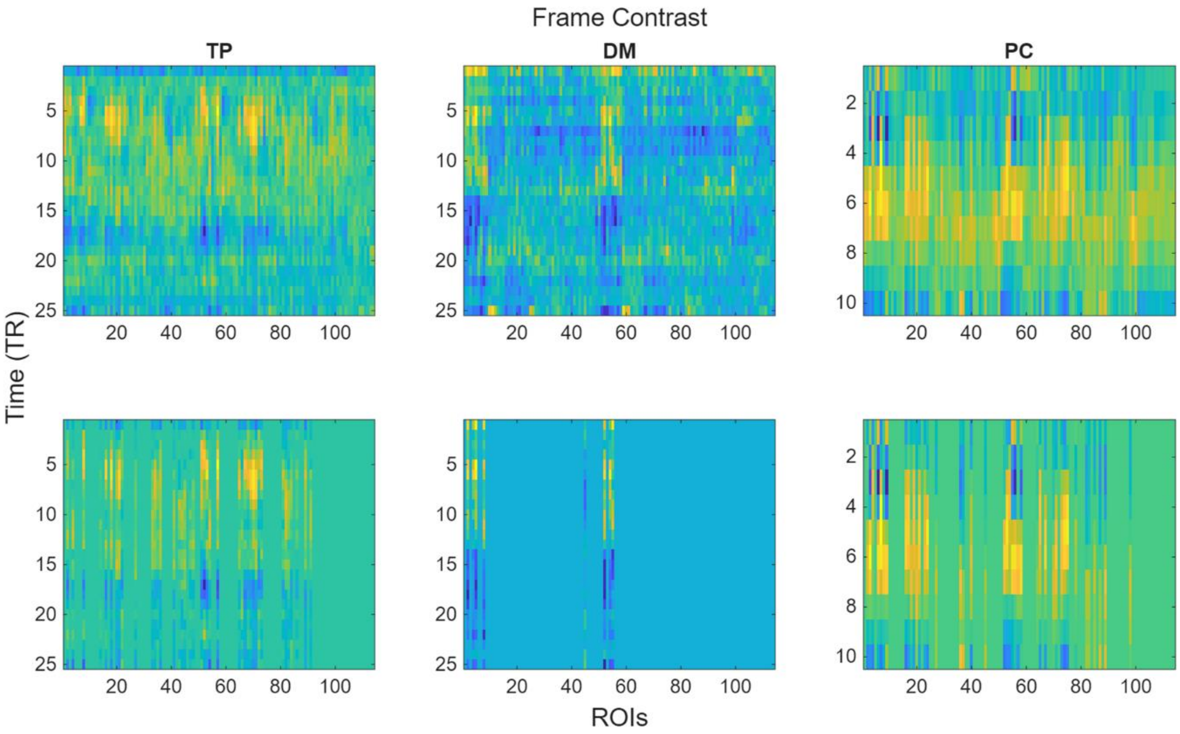

**Supplementary Figure S4. FIR deconvolution profiles for auditory pitch.** Panels illustrate the estimated system response functions across 114 regions of interest (ROIs) for the movies *The Present* (TP, left) and *Despicable Me* (DM, right). **Top row:** Unthresholded mean FIR response profiles across all 114 ROIs. **Bottom row:** Significant regional response functions identified through group-level statistical inference. Statistical significance was determined by aggregating individual-level *F*-tests using Fisher's method, with the resulting group-level maps thresholded at a false discovery rate (FDR) of $p < 0.05$.

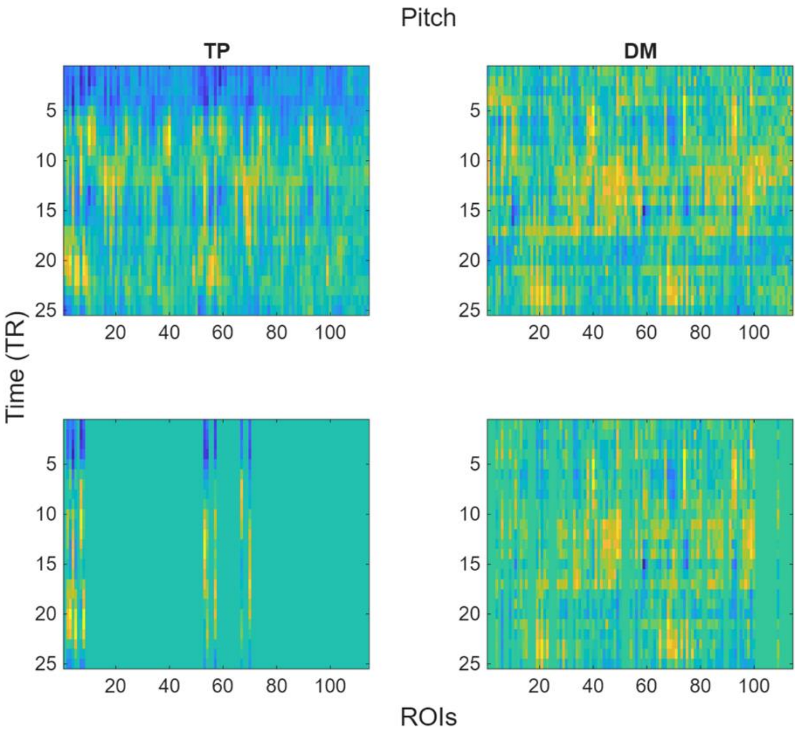

**Supplementary Figure S5. Temporal orchestration of frame luminance, pupillary dynamics, and system response functions. (A)** Continuous time series of the group-median pupil size and frame luminance (mean pixel intensity) during the movie *The Present*. **(B)** Cross-correlation function between frame luminance and pupil size, demonstrating a prominent negative peak at approximately 1.0 second, reflecting the intrinsic biological latency of the pupillary light reflex. A positive lag indicates that luminance fluctuations precede the pupillary response (*Luminance lead*), while a negative lag indicates that the pupillary signal precedes luminance changes (*Pupil size lead*). **(C)** Empirical Physiological Response Function (PRF) for the pupil, estimated using frame luminance as the input vector. A canonical hemodynamic response function (HRF) is overlaid to demonstrate the distinct timescales and sequential temporal relationship between the two profiles. While the upstream autonomic PRF peaks early (~1.0 s), the downstream neurovascular HRF peaks significantly later (~5–6 s), illustrating the chronological temporal cascade from peripheral physiological tracking to localized cerebral blood flow.

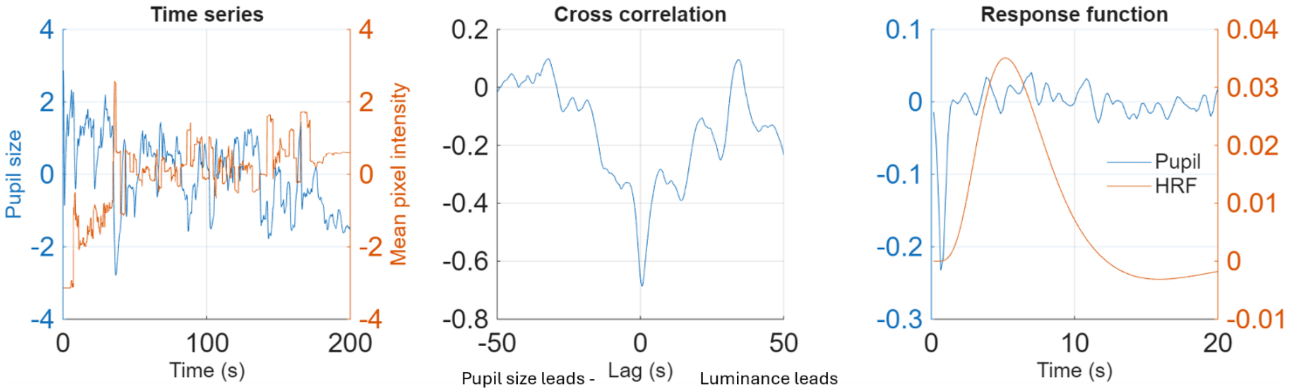

**Supplementary Figure S6. FIR deconvolution profiles for pupil size.** Panels illustrate the estimated system response functions across 114 regions of interest (ROIs) for the movie *The Present* (TP). **Left panel:** Unthresholded mean FIR response profiles across all 114 ROIs. **Right panel:** Significant regional response functions identified through group-level statistical inference. Statistical significance was determined by aggregating individual-level *F*-tests using Fisher's method, with the resulting group-level maps thresholded at a false discovery rate (FDR) of $p < 0.05$.

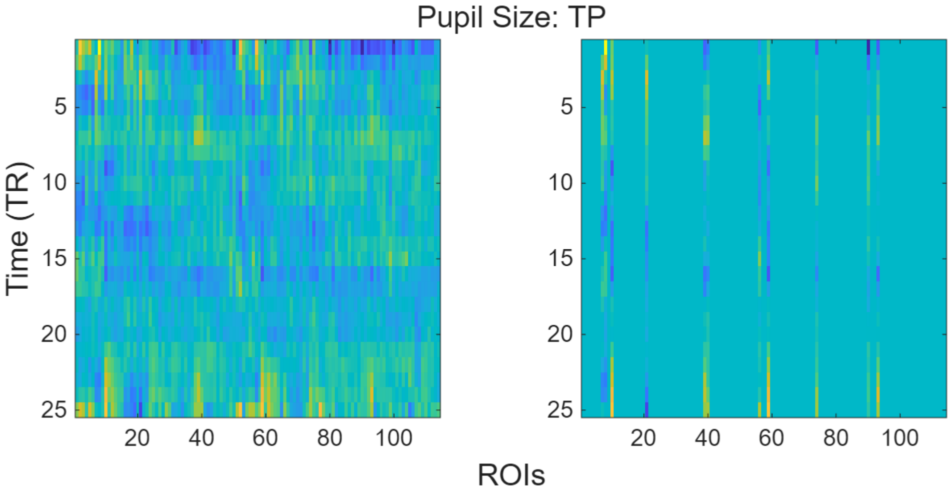

**Supplementary Figure S7. Spatiotemporal cross-correlation matrices for convolved and raw Theory of Mind (ToM) ratings.** Matrices illustrate the relationship between fMRI activity across 114 regions of interest (ROIs) and ToM ratings during the movies *The Present* (TP, first column), *Despicable Me* (DM, second column), and *Partly Cloudy* (PC, third column). The top row displays correlations using canonical HRF-convolved ToM ratings, while the bottom row displays correlations using the raw (unconvolved) ToM ratings. A positive lag indicates that the stimulus/feature time series precedes the BOLD signal (i.e., Feature lead), whereas a negative lag indicates that the BOLD signal precedes the feature (i.e., BOLD lead). Vertical dividers delineate ROI assignments across the left and right hemispheres according to Yeo's seven-network parcellation, as well as subcortical regions. The comparison highlights how utilizing the raw behavioral trace mitigates the "double-smoothing" artifact, resulting in more physiologically plausible temporal alignments.

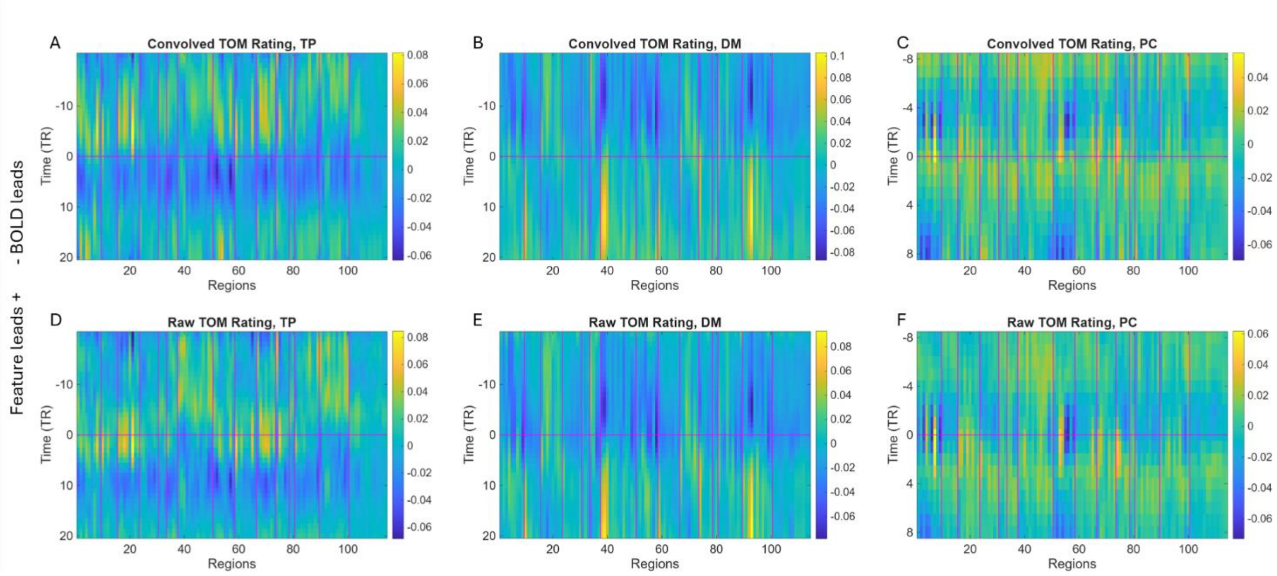

**Supplementary Figure S8. FIR deconvolution profiles for theory of mind (ToM) rating.** Panels illustrate the estimated system response functions across 114 regions of interest (ROIs) for the movies *The Present* (TP, first column), *Despicable Me* (DM, second column), and *Partly Cloudy* (PC, third column). **Top row:** Unthresholded mean FIR response profiles across all 114 ROIs. **Bottom row:** Significant regional response functions identified through group-level statistical inference. Statistical significance was determined by aggregating individual-level *F*-tests using Fisher's method, with the resulting group-level maps thresholded at a false discovery rate (FDR) of $p < 0.05$.

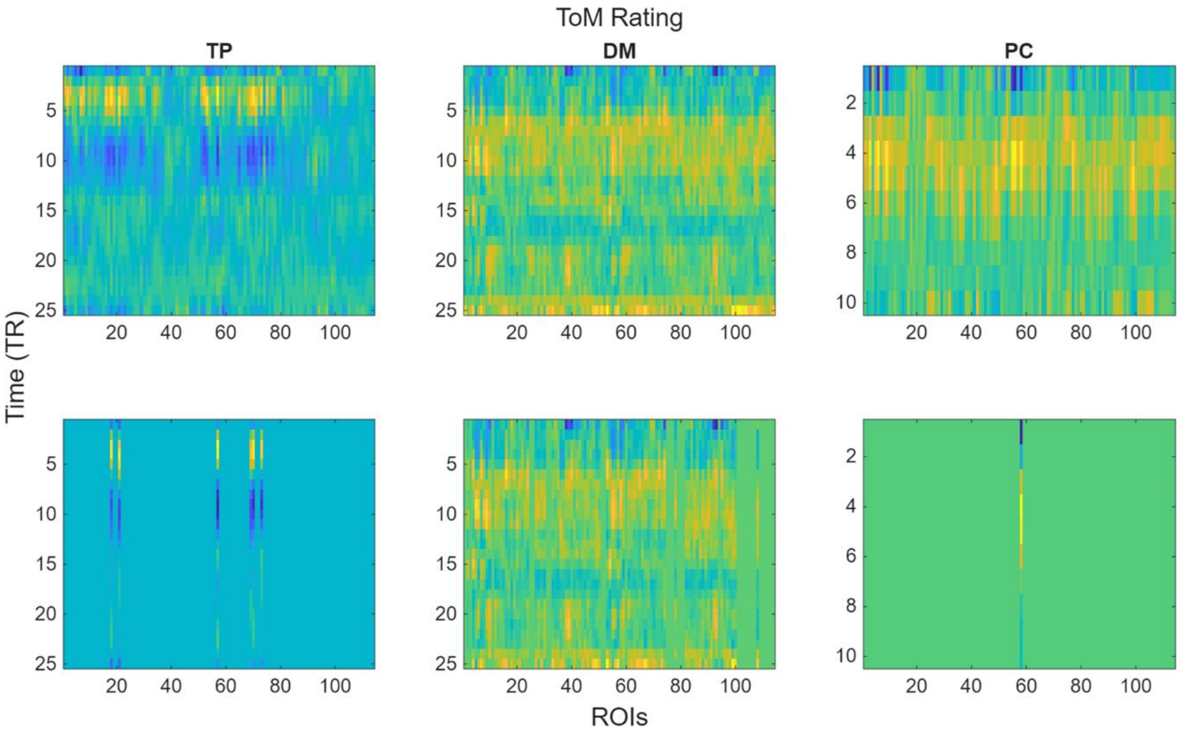

**Supplementary Figure S9. Cross-individual split-half consistency of finite impulse responses (FIR) across all features and movie conditions.** Bar plots illustrate the temporal correlation (r) between group-averaged FIR profiles from independent split-half cohorts across all extracted stimulus features and the three movie datasets (TP: *The Present*; DM: *Despicable Me*; PC: *Partly Cloudy*). Vertical dividers delineate region of interest (ROI) assignments across the left and right hemispheres according to Yeo's seven-network parcellation, alongside subcortical structures. This comprehensive mapping highlights a consistent sensory-to-associative gradient in response shape reliability across different feature classes.

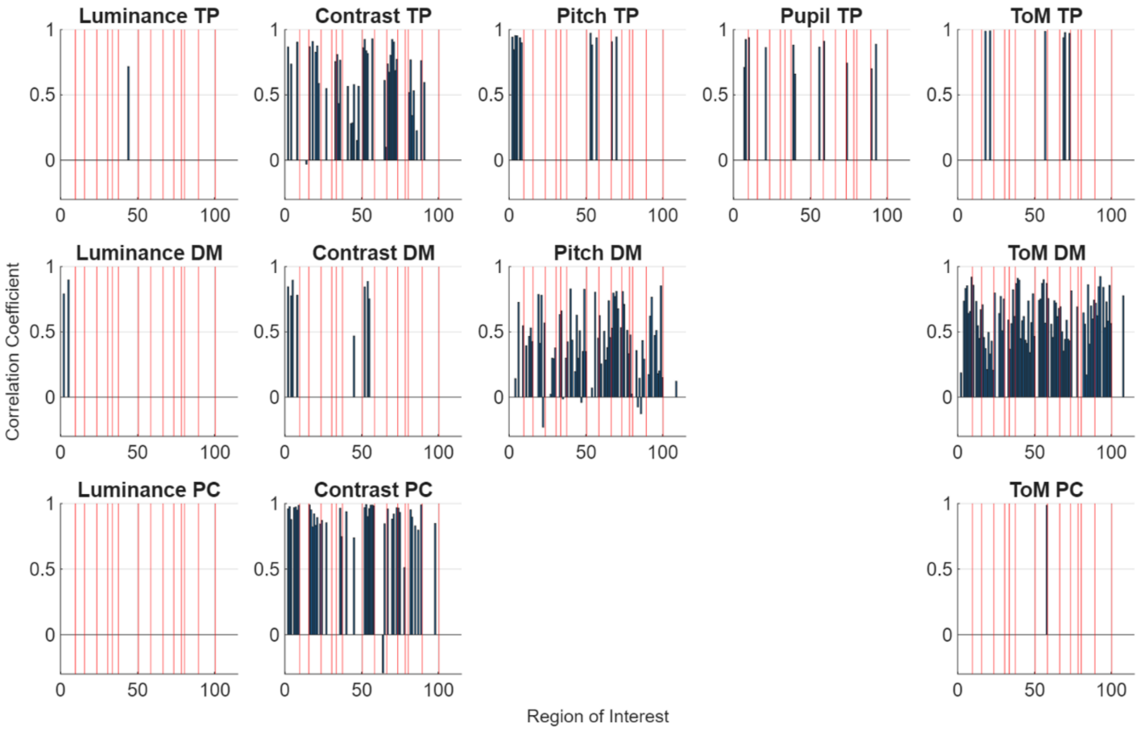

**Supplementary Figure S10. Cross-individual sign consistency of finite impulse responses (FIR) across all features and movie conditions.** Bar plots illustrate the cross-subject sign consistency of estimated FIR profiles across all stimulus features and the three movie datasets (TP: *The Present*; DM: *Despicable Me*; PC: *Partly Cloudy*). Values represent the proportion of participants sharing the same response polarity (positive versus negative BOLD fluctuations) relative to the dominant group sign. Vertical dividers delineate ROI assignments across the left and right hemispheres according to Yeo's seven-network parcellation, as well as subcortical structures. Note that the baseline chance level for sign consistency is 0.5.

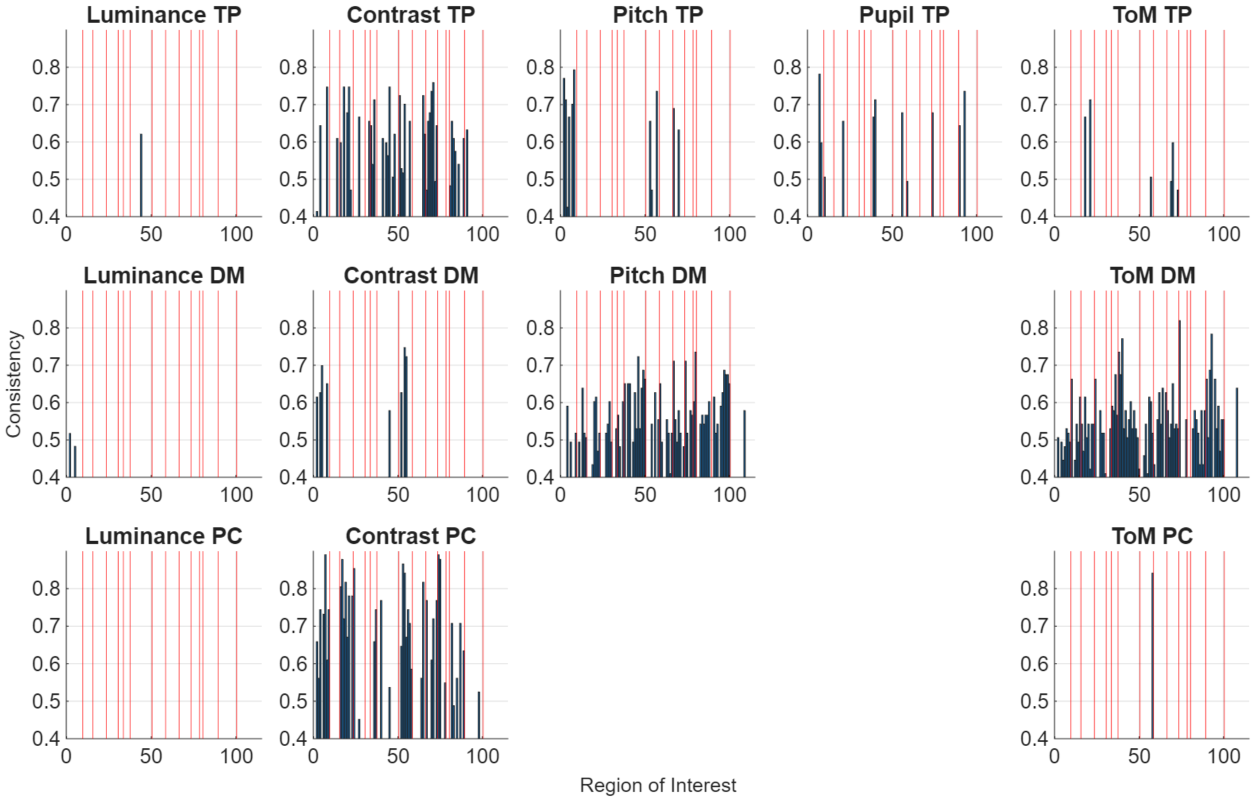
